# Molecular basis of chemosensing by internal sensory neurons in *Drosophila* larvae

**DOI:** 10.1101/2024.01.20.576342

**Authors:** Shan Lu, Cheng Sam Qian, Amena Khair-Eldin, Wesley B. Grueber

**Affiliations:** Zuckerman Mind Brain Behavior Institute, Jerome L. Greene Science Center, 3227 Broadway, L9-007, Columbia University, New York, NY 10027; Department of Biological Sciences, Jerome L. Greene Science Center, 3227 Broadway, L9-007, Columbia University, New York, NY 10027; Department of Physiology and Cellular Biophysics, Jerome L. Greene Science Center, 3227 Broadway, L9-007, Columbia University, New York, NY 10027; Department of Neuroscience, Jerome L. Greene Science Center, 3227 Broadway, L9-007, Columbia University, New York, NY 10027

## Abstract

Interoceptive neurons monitor the chemical and physical state of the body, providing critical information to the central nervous system that maintains homeostasis and promotes survival. The circuit and molecular basis for these interoceptive senses remain under active investigation. A population of larval *Drosophila* sensory neurons, the tracheal dendrite (td) neurons, extend dendrites along respiratory organs and provide a tractable model for investigating the cellular and molecular basis of internal chemosensation. Here, we show that partially overlapping populations of td neurons respond to decreases in O_2_ and increases in CO_2_. To define the molecular basis of these responses, we examined the roles of atypical soluble guanylyl cyclases (Gycs) and gustatory receptors (Grs). We found that Gyc88E/Gyc89Db were required for hypoxia-evoked activity, whereas Gr28b was required for responses to CO_2_. Targeted expression of Gr28b.c isoform in td neurons restored responses to CO_2_ in mutant larvae and conferred ectopic sensitivity to CO_2_ within the td network. We further found that gas-sensitive td neurons were activated upon prolonged larval burrowing, a natural-like feeding-associated behavior. Together, these findings reveal molecular mechanisms that enable interoceptive neurons to monitor respiratory gases and advance a *Drosophila* model for studying the neural and molecular basis of internal chemosensation.

## INTRODUCTION

Interoceptive neurons detect physiological states within the body and enable homeostasis and adaptive behavior. Across species, these neurons innervate visceral organs and relay information about their chemical and physical state to the central nervous system (CNS). In mammals, internal sensory neurons respond to diverse internal cues, including nutrients, temperature, mechanical stretch, irritants, and hormones (Fajardo et al., 2008; Hillsley & Grundy, 1998; Paintal, 1973; Williams et al., 2016). Despite extensive characterization, many of the molecular mechanisms that allow interoceptive neurons to detect and encode these signals remain poorly understood. The genetically accessible nervous system in *Drosophila* larvae offers a powerful and tractable model for defining the cellular and molecular logic of internal sensory transduction.

The tracheal dendrite (td) interoceptive neurons of larval *Drosophila* project dendrites across the tracheal (respiratory) system in a segmental pattern. Of the 13 bilateral td neurons, nine project axons to the subesophageal zone (SEZ), a chemosensory recipient region in the brain, whereas others project to a thoracic region of the ventral nerve cord (VNC) of unknown significance (Qian et al., 2018). All td neurons express at least one reporter in the gustatory receptor (GR) or ionotropic receptor (IR) families (Qian et al., 2018), which, together with olfactory receptors (ORs), make up the three major chemoreceptor families in *Drosophila* (Joseph & Carlson, 2015). In addition, a subset of td neurons co-expresses the atypical soluble guanylyl cyclases Gyc88E and Gyc89Db, which respond to decreases in O_2_ levels (Langlais et al., 2004). Roles for GRs, IRs, and Gycs in td neurons are not known; however, extensive molecular and anatomical heterogeneity of td neurons raise the possibility of diverse functions in interoception (Qian et al., 2018).

Based on their dendritic projections along specific tracheal branches, td neurons were initially proposed to function as gas sensors or proprioceptors (Bodmer & Jan, 1987). Two recent studies have begun to shed light on possible sensory roles in nociception and CO_2_ sensing (Huckesfeld et al., 2021; Imambocus et al., 2022). In the latter study, axon projections to the SEZ showed calcium responses upon animal exposure to high environmental CO_2_ (Huckesfeld et al., 2021). The specific identity of CO_2_ responsive td neurons and molecules underlying these responses are still open questions. The molecular basis of CO_2_-sensing has been previously investigated in the olfactory system of adult and larval *Drosophila* (Faucher et al., 2006; Kwon et al., 2007). Here, CO_2_ is sensed by two receptors, Gr21a and Gr63a, which are co-expressed in sensory neurons that innervate specific sensilla on the body wall of both larvae and adults (Jones et al., 2007; Kwon et al.). In addition, a population of gustatory neurons respond to aqueous and gaseous CO_2_ (Fischler et al., 2007), and the responses require Ir25a, Ir56d and Ir76b (Sanchez-Alcaniz et al., 2018). Whether additional GRs, IRs, other molecules, or some combination of the above known players mediate CO_2_ detection in td neurons is yet to be investigated.

Here, we study the responses of td neurons to changes in O_2_ and CO_2_ levels using the calcium integrator CaMPARI2 (Fosque et al., 2015). Our results show that specific and partially overlapping subsets of td neurons are responsive to changes in external O_2_ and CO_2_ levels, as well as in a naturalistic condition in which gas exchange is impaired. We used genetic loss- and gain-of-function experiments to identify molecules that mediate td responses to changes in O_2_ and CO_2_ levels. We provide evidence that Gycs mediate the response of td neurons to O_2_ and that Gr28b.c mediates responses to CO_2_. We propose that this subset of td neurons functions to monitor internal gas levels and that other td neurons could detect a range of other interoceptive signals in *Drosophila* larvae.

## RESULTS

### td neurons associate with the basal surface of tracheal epithelial cells

Larval td neurons extend their dendrites primarily along a specific tracheal branch called the lateral trunk. The dendrites associate closely with tracheal epithelial cells (Qian et al., 2018) (***Figure 1A-B***). To further characterize the association between td dendrites and the lateral tracheal trunk, we imaged td dendrites, tracheal epithelial cells, and autofluorescent tracheal cuticle (***Figure 1C***). Orthogonal views showed that td dendrites associate closely with epithelial cells but at the level of resolution achieved (approximately 0.1μm), revealed no evidence for dendrite exposure to the tracheal lumen (***Figure 1C_1_-C_5_***) (Qian et al., 2018).

**Figure 1.**
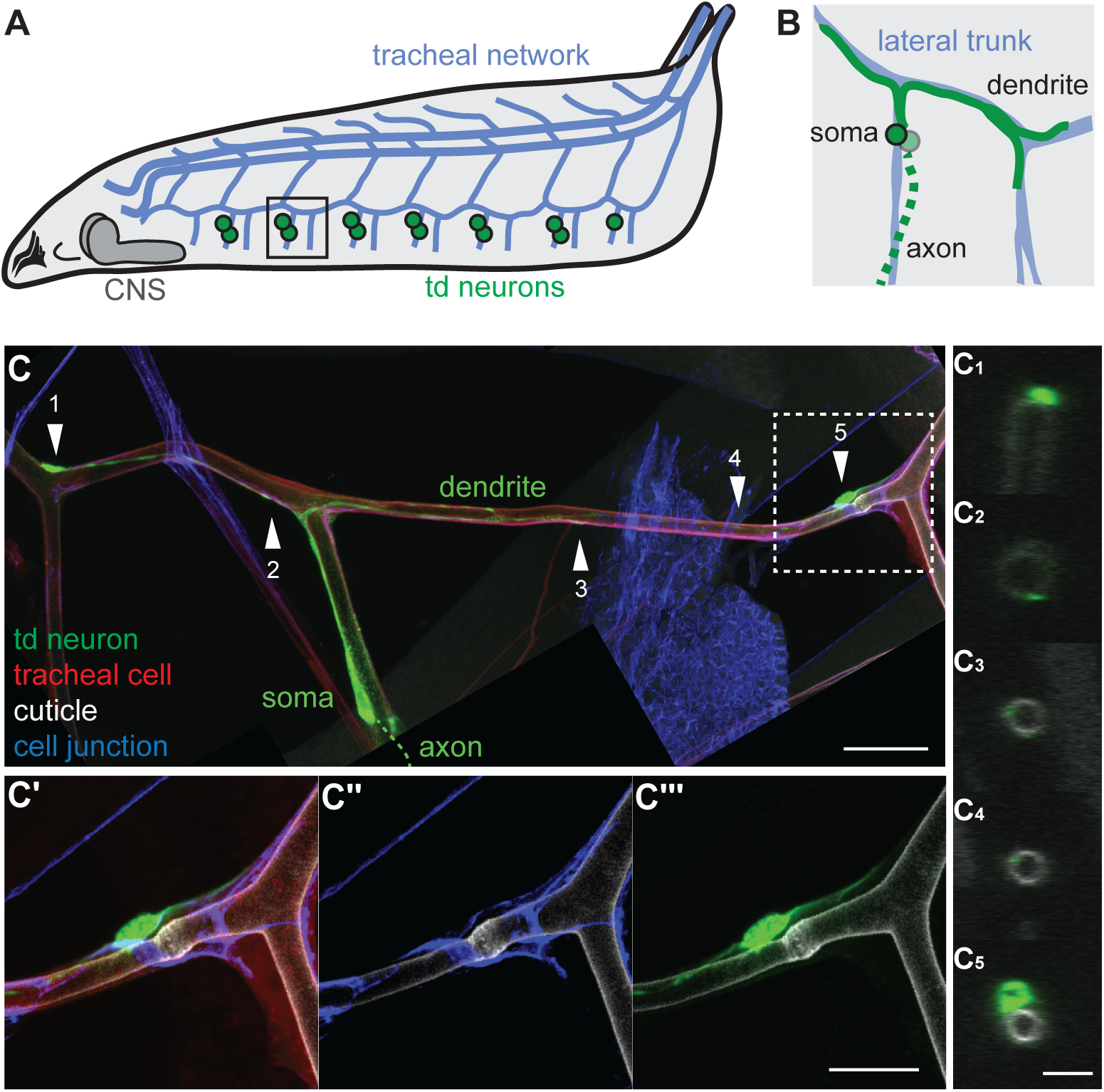
Characterization of td neuron associations with tracheae. **A.** Schematic of *Drosophila* larva and td neuron cell bodies (green circles) in relation to major tracheal network (blue lines). **B.** Schematic of one pair of td neurons in the black box from **A**, showing cell bodies (green circles), dendrite (solid green lines), and axon (dashed green line). td dendrites are closely associated with specific tracheal branches (solid blue lines). Most of td dendrite length extend along the lateral trunk. **C.** Image of td neuron (green, *Gr28a-QF2^G4H^*), tracheal epithelial cells (red, *breathless-Gal4*), tracheal cuticle (tracheal lumen surrounded by cuticle, grey, autofluorescence), and tracheal cell junctions (blue, anti-Coracle) labelled together. td dendrite is closely associated with tracheae. **C1-C5.** Orthogonal views of td dendrite in relation to tracheal cuticle as indicated by corresponding arrowheads in **C**. **C’**, Zoom-in image of the dashed line box in **C**. **C’’**, Tracheal lumen forms “donut-shaped” enlargement (indicated by the arrowhead #5) at the fusion site guided by tracheal fusion cells. The enlargement shows stronger autofluorescence under 405nm laser. **C’’’**, td dendrite forms enlargement near tracheal fusion site. **A** and **B** adapted from (Qian et al., 2018) and modified with permission. Scale bars: **C, D,** 20μm; **C’-C’’’,** 10μm; **C1-C5,** 5μm.

We also tested for td dendrite embedding vs. surface localization using a Triton X-100-free immunohistochemistry procedure that occludes HRP labeling in regions of neurons that are surrounded by glia or embedded in epithelial cells (Kim et al., 2012) (***Figure 1 – figure supplement 1A-B***). The procedure led to reduced HRP labelling of td cell bodies, likely due to glial ensheathment, but provided no evidence for enclosure of td dendrites by tracheal epithelial cells (***Figure 1 – figure supplement 1C***). We conclude that td sensory dendrites are closely apposed to tracheal epithelial cells and are directly exposed to hemolymph along most, if not all, of their length.

### td neurons detect decreases in O_2_ level

The locations of td neurons along respiratory organs led to the hypothesis that they function in mechanosensation or chemosensation (Bodmer & Jan, 1987). The expression of a variety of gustatory receptor (GR) and ionotropic receptor (IR) reporters in td neurons, as well as their axon projections, are more consistent with chemosensory roles for at least a subset of td neurons (Qian et al., 2018). We nevertheless checked for expression of mechanosensory receptors NompC and Piezo using Gal4 reporter lines but found no expression in td neurons (***Figure 2 – figure supplement 1A-B***). Likewise, previous studies of expression *tmc-Gal4* reported expression in class I, class II, and bd neurons, but described no characteristic td neuron axon projections in the larval CNS (Guo et al., 2016).

**Figure 2.**
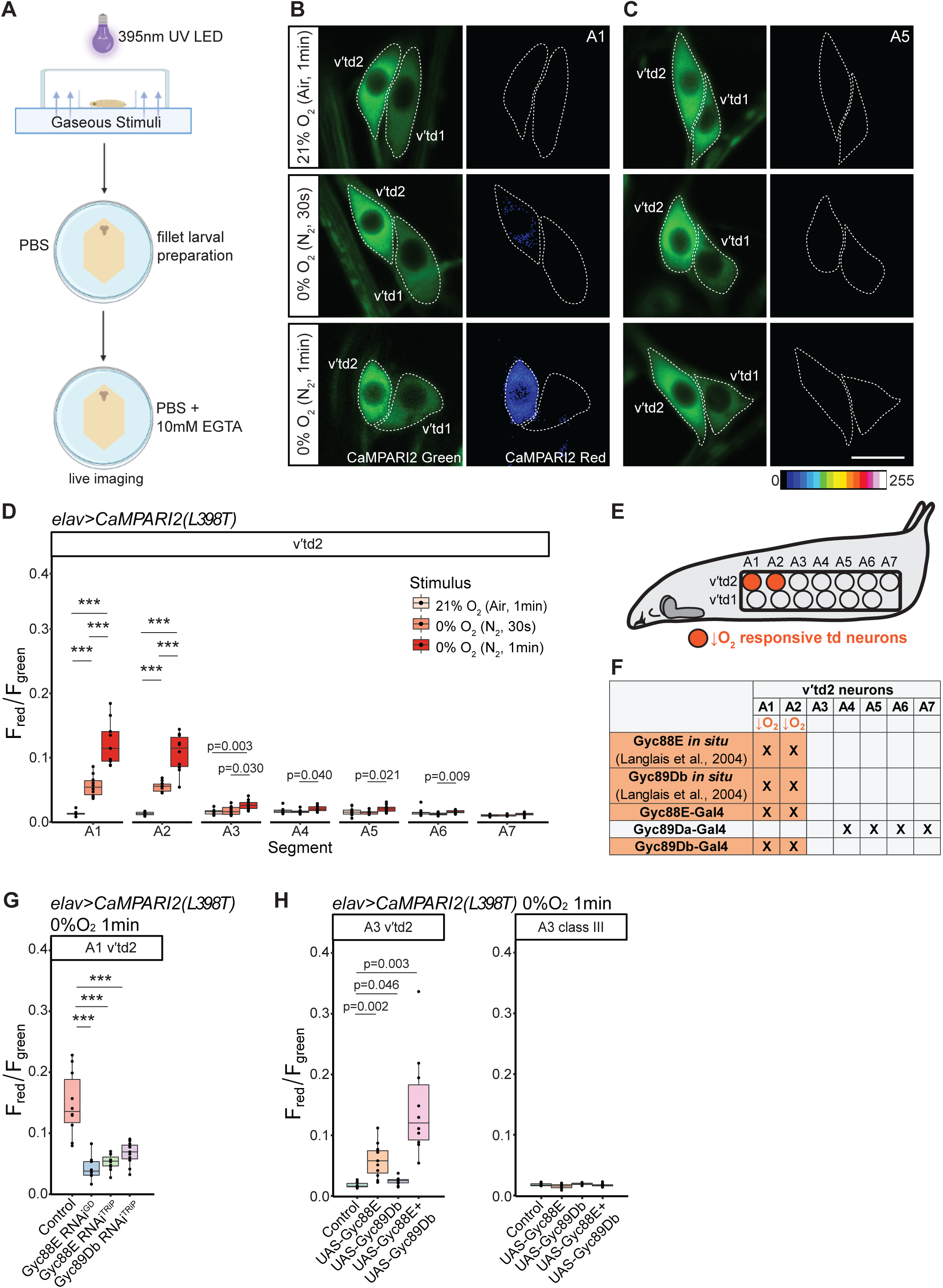
td neurons detect decreases in O2 level. **A.** Schematic of gas-delivery and post-hoc neural imaging of CaMPARI2 responses to gaseous stimuli. (Created with BioRender.com). **B-C.** Green and red CaMPARI2 fluorescence in A1 and A5 td neurons, respectively, in response to decreases in O2 level. (Red channel shown in pseudo color for better visualization). Dorsal is top, and anterior is to the left. Larvae exposed to 1 minute of air (21% O2) and UV showed little to no green to red CaMPARI2 photoconversion in A1 and A5 td neurons. Larvae exposed to 30 seconds or 1minute of hypoxia (0% O2) and UV showed increased green to red photoconversion in A1 v’td2 neuron but not in A1 v’td1 neuron or A5 td neurons. CaMPARI2 was expressed pan-neuronally (*elav-Gal4>UAS-CaMPARI2(L398T)*). **D.** Quantification of red/green CaMPARI2 fluorescence in all v’td2 neurons across segments A1-A7 in response to hypoxia. A1-A2 v’td2 neurons responded to 30 seconds or 1 minute of hypoxia. n=9-10 animals per condition. **E.** Schematic of the relative positions of the two hypoxia-sensitive td neurons on one side of the larval body wall. Schematic of larva adapted from (Qian et al., 2018) and modified with permission. **F.** Hypoxia-sensitive td neurons express distinct Gycs. Orange text indicates stimulus detected by td neurons. Following rows with crosses indicate Gyc expression patterns of td neurons. Gyc *in situ* hybridizations were performed in embryonic stages (Langlais et al., 2004)). **G.** Quantification of red/green CaMPARI2 fluorescence in the hypoxia-sensitive A1 v’td2 neuron in control and loss-of-function conditions in response to 1 minute of 0% O2. RNAi knockdown of Gyc88E or Gyc89Db under the control of *elav-Gal4* significantly decreased the response to hypoxia. n=10-11 animals per condition. **H.** Quantification of red/green CaMPARI2 fluorescence in the hypoxia-insensitive A3 v’td2 neuron and A3 class III somatosensory neuron in control and gain-of-function conditions in response to 1 minute of 0% O2. Overexpression of Gyc88E alone or Gyc88E together with Gyc89Db resulted in significant increases in fluorescence ratio in A3 v’td2 neurons in response to hypoxia. Overexpression of Gycs did not induce response to hypoxia in A3 class III somatosensory neuron. n=9-11 animals per condition. Scale Bar, 10μm in all panels.

Prior studies found expression of the molecular O_2_ sensors guanylyl cyclase (Gyc)88E and Gyc89Db in a subset of td neurons and showed that Gycs detect a decline in O_2_ levels *in vitro* (Langlais et al., 2004). We used the genetically encoded calcium integrator CaMPARI2, which undergoes permanent green-to-red photoconversion in the presence of high intracellular calcium and UV light, to test whether respiratory gases could stimulate td neurons. Specifically, we used CaMPARI2(L398T) which is the low affinity version of the CaMPARI2 molecule (Fosque et al., 2015; Moeyaert et al., 2018). We placed third-instar larvae in a chamber and exposed them to 21% O_2_ (79% N_2_, 0.04% CO_2_) or 0% O_2_ (100% inert N_2_). During 30 seconds or 1 minute gas delivery we simultaneously applied UV light to enable CaMPARI2 photoconversion, ensuring that CaMPARI2 only captured Ca^2+^ increases during this time window (***Figure 2A***). We then dissected the larvae and imaged red and green fluorescence in all 13 segmental td neurons for post hoc assessment of neural calcium responses during gas delivery. When larvae were exposed to air, td neurons showed virtually no green-to-red photoconversion (***Figure 2B-C***), suggesting that td neurons are largely inactive in normoxic conditions and are not responsive to UV light in this assay. However, when larvae were exposed to 0% O_2_ the A1 and A2 v′td2 neurons showed significantly higher red/green fluorescence ratio compared to controls after 30s and 1 min exposure (***Figure 2B-C***), suggesting that these two td neurons are activated by O_2_ decreases (***Figure 2D-E***). Other td neurons showed modest responses to 1 minute exposure to 0% O_2_, potentially due to perturbation of normal internal homeostatic states induced by prolonged hypoxic condition (***Figure 2D and Figure 2 - figure supplement 1D***). We note that, because td neurons are internal to the body, the concentration of gases experienced by td neurons are likely different from the concentrations that we delivered externally.

### Guanylyl cyclases mediate responses of td neurons to low oxygen

Based on differential sensitivities to O_2_ depletion, we classified A1-A2 v’td2 neurons as oxygen-sensitive (***Figure 2E***). Prior studies reported expression of guanylyl cyclases Gyc88E and Gyc89Db RNA in td neurons (Langlais et al., 2004). We confirmed expression using Gal4 reporter lines and additionally found that a Gyc89Da-Gal4 reporter line labels a subset of abdominal td neurons (***Figure 2F* and *Figure 2 – figure supplement 2A-C***). To determine whether Gycs mediate responses to 0% O_2_, we used RNAi to knock down Gyc88E or Gyc89Db, and measured td responses in larvae exposed to 0% O_2_ for 1 minute. Since A1 and A2 td neurons behave similarly, we used the A1 v’td2 neuron as a representative cell. Indeed, we found that knockdown of Gyc88E or Gyc89Db significantly decreased responses of v’td2 to 0% O_2_ (**Figure 2G**).

To determine if Gyc88E or Gyc89Db alone, or in combination, are sufficient for O_2_ responses, we misexpressed these subunits in td neurons and measured CaMPARI2 responses in animals exposed to 0% O_2_. We monitored responses in A3 neurons (both v’td2 and v’td1) as they do not normally show detectable expression of either Gyc subunit (***Figure 2F and Figure 2 – figure supplement 2***). We found that ectopic expression of Gyc88E and Gyc89Db together enhanced responses to 0% external O_2_ in A3 td neurons (***Figure 2H and Figure 2 – figure supplement 2E***). Ectopic expression of Gyc88E alone was sufficient to induce weaker responses to 0% O_2_, whereas ectopically expressing Gyc89Db alone induced no such response (**Figure 2H *and Figure 2 – figure supplement 2E***). A class III peripheral somatosensory neuron did not acquire sensitivity to hypoxia upon co-expression of Gyc88E and Gyc89Db (***Figure 2H***). Together, our results are consistent with previous *in vitro* studies and support a role for Gycs in responses of td neurons to decreases in O_2_. Furthermore, tracheal association appears to be important for these responses.

### td neurons detect increases in CO_2_ level

Prior studies showed calcium responses to high CO_2_ in axon bundles of td neurons leading to the SEZ, but the specific identity of cells showing these responses was not addressed (Huckesfeld et al., 2021; Qian et al., 2018). To determine the identity of neurons that detect CO_2_ we exposed larvae to different concentrations of CO_2_ for 2 minutes and quantified CaMPARI2 responses. CaMPARI2 was driven by pan-neural *elav-Gal4* in these experiments to capture responses in all td neurons. We found that 10-20% CO_2_ (with O_2_ held constant at atmospheric 20-21% levels and balanced with inert N_2_) elicited responses in A1, A2, and A3 td neurons (***Figure 3A-B and Figure 3 – figure supplement 1***), indicating partial overlap in the neurons that detect decreases in O_2_ and increases in CO_2_. By contrast, the A3 td showed only weak sensitivity to low O_2_ and responded strongly to high CO_2_ levels, indicating specificity for CO_2_. We exposed larvae to 79-80% and 100% CO_2_ for 30 seconds (***Figure 3A and C***), or 79-80% CO_2_ for 15 seconds (***Figure 3 – figure supplement 1B***), and detected responses in the same three neurons. We conclude that td neurons show concentration-dependent responses to increased CO_2_ levels.

**Figure 3.**
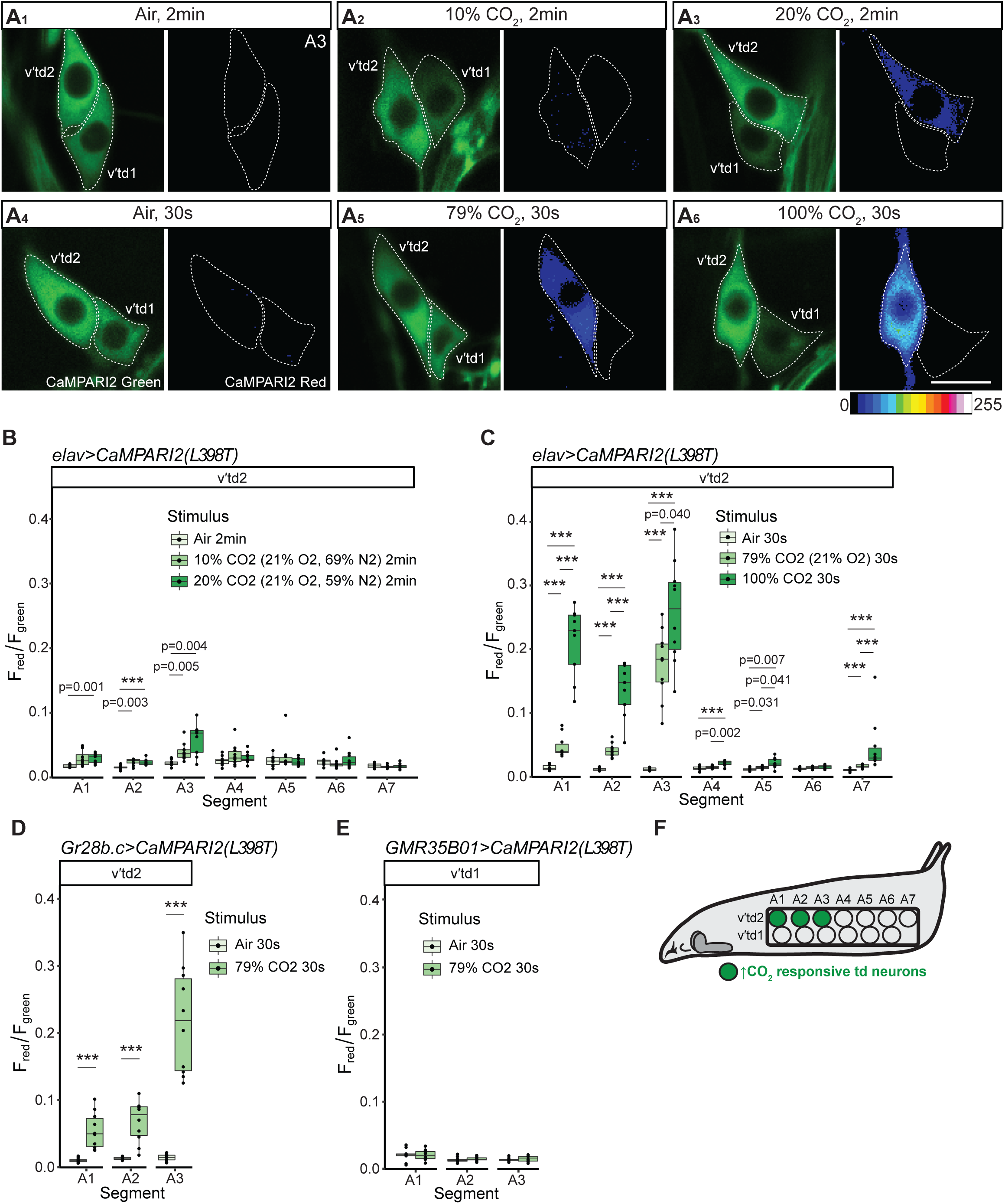
td neurons detect increases in CO2 level. **A.** Green and red CaMPARI2 fluorescence in A3 td neurons in response to increases in CO2 level. (Red channel shown in pseudo color). Dorsal is top, and anterior is to the left. **A1, A4.** Larvae exposed to 2 minutes or 30 seconds of air (∼0.04% CO2) and UV showed no green to red CaMPARI2 photoconversion in A3 td neurons. **A2-3, A5-6.** Larvae exposed to 2 minutes of 10% CO2, 20% CO2 (21% O2, balanced with N2), or 30 seconds of 79% CO2 (21% O2), 100% CO2 and UV showed increased green to red photoconversion in A3 v’td2 neuron but not in v’td1 neuron. **B-C.** Quantification of red/green CaMPARI2 fluorescence in all v’td2 neurons across segments A1-A7 in response to different CO2 levels. CaMPARI2 expression was driven by *elav-Gal4*. A1-A3 v’td2 neurons responded to 2 minutes of 10% CO2, 20% CO2, and 30 seconds of 79% CO2, 100% CO2. n=8-10 animals per condition. **D-E.** Quantification of red/green CaMPARI2 fluorescence in A1-A3 td neurons in response to 79% CO2. CaMPARI2 expression was driven by *Gr28b.c-Gal4* for v’td2, and *GMR35B01-Gal4* for v’td1. Same three td neurons (A1-A3 v’td2) responded to 79% CO2. n=9-10 animals per condition. **F.** Schematic of the relative positions of the three CO2-sensitive td neurons on one side of the larval body wall. Schematic of larva adapted from (Qian et al., 2018) and modified with permission. Scale Bar, 10μm in all panels.

To confirm the identity of td neurons that responded to increases in CO_2_, we repeated these assays with v’td2 and v’td1 specific drivers, *Gr28b.c-Gal4* **(*Figure 3 – figure supplement 1C*)** and *GMR35B01-Gal4* **(*Figure 3 – figure supplement 1D*)**, respectively, to drive CaMPARI2 expression (Qian et al., 2018). We observed that responses to CO_2_ in segments A1-A3 were specific to the v’td2 neuron (***Figure 3D-F***). These data indicate that subsets of td neurons are sensitive to both O_2_ and to CO_2_, with one td neuron selectively sensitive to CO_2_.

### Gr28b is necessary for responses to CO_2_ in td neurons

To investigate the molecular basis of CO_2_ sensing we focused on A3 v’td2 responses since this cell showed robust responses to CO_2_ but comparatively weaker responses to hypoxia. We previously reported expression of GR-GAL4 and IR-GAL4 reporters in td neurons (Qian et al., 2018). The functional significance of the specific complement of GRs and IRs expressed in td neurons remains unknown. td neurons appear to lack expression of the olfactory CO_2_-receptor combination, Gr21a/63a, as the reporter lines for these two GRs showed no distinctive td-like axon projections in the larval CNS (Kwon et al., 2011). A3 v’td2 neurons also do not express reporters for all components of the gustatory carbonation-sensing receptor combination, Ir25a/56d/76b, as expression of *Ir25a-Gal4* and *Ir76b-Gal4* is lacking (Qian et al., 2018) (***Figure 3 – figure supplement 1E-F***), and Ir56d is not reported to be expressed in the larval stage (Sanchez-Alcaniz et al., 2018; Stewart et al., 2015). These studies suggest there are new mediators for td responses to CO_2_. The observation that CO_2_-sensitive v’td2 neurons express a unique combination of Gr28a, Gr28b.c, and Gr89a (***Figure 4A***) provided candidates to test for roles in CO_2_ detection (Qian et al., 2018).

**Figure 4:**
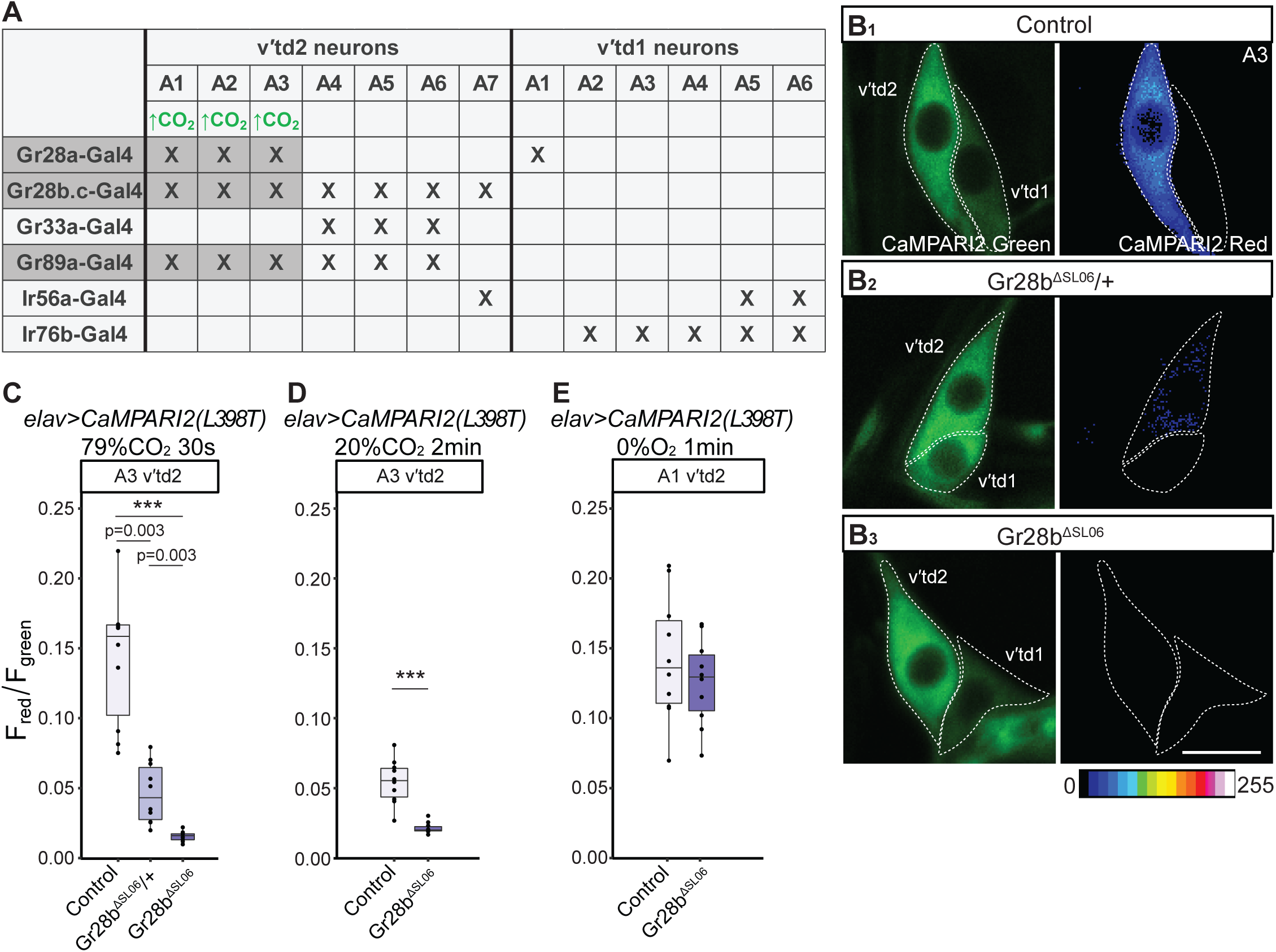
Gr28b is necessary for responses to CO2 in td neurons. **A.** Gas-sensitive td neurons express distinct chemosensory receptors. Green text indicates stimulus detected by td neurons. Following rows with crosses indicate chemosensory receptor reporter expression patterns of td neurons. Data from (Qian et al., 2018). Gray boxes highlight GR reporters that are expressed in CO2-sensitive td neurons characterized in Figure 3. The combined reporter expression of Gr28a, Gr28b.c, and Gr89a is unique to CO2-sensitive td neurons. **B.** Green and red CaMPARI2 fluorescence in A3 td neurons in control (**B1**), *Gr28b^ΔSL06^* heterozygous (**B2**), and *Gr28b^ΔSL06^* (**B3**) larvae in response to 79% CO2. (Red channel shown in pseudo color). Dorsal is top, and anterior is to the left. **B1.** Control larvae exposed to 79% CO2 showed green to red CaMPARI2 photoconversion in A3 v’td2 neuron (as in **Figure 3A5**). **B2-3.** *Gr28b^ΔSL06^* heterozygous, and *Gr28b^ΔSL06^*larvae exposed to 79% CO2 showed decreased, and no detectable green to red photoconversion, respectively, in A3 v’td2 neuron. **C-D.** Quantification of red/green CaMPARI2 fluorescence in A3 CO2-sensitive v’td2 neuron in control and Gr28b mutant animals in response to 30 seconds of 79% CO2 and 2 minutes of 20% CO2. *Gr28b^ΔSL06^*heterozygous, and *Gr28b^ΔSL06^* animals showed decreased responses to 79% CO2. *Gr28b^ΔSL06^* animals also showed decreased responses to 20% CO2. n=10 animals per condition. **E.** Quantification of red/green CaMPARI2 fluorescence in A1 hypoxia-sensitive v’td2 neuron in control and Gr28b mutant animals in response to 1 minute of 0% O2. *Gr28b^ΔSL06^* animals showed normal responses to hypoxia compared to control animals. n=10 animals per condition. Scale Bar, 10μm in all panels.

As a first test of whether Gr28a, Gr28b, or Gr89a are necessary for td neuron responses to CO_2_, we measured the responses to 30 seconds of 100% CO_2_ in larvae with reduction or loss of GR function. We used RNAi or, where available, mutant lines to knock down or knock out specific Grs and examined CaMPARI2 responses in A3 v’td2 neurons. We found that *Gr28a^1^* mutants and Gr89a RNAi knockdown animals showed normal responses to 100% CO_2_. However, Gr28b RNAi knockdown significantly reduced responses to 100% CO_2_ (***Figure 4 – figure supplement 1***). We observed a similar phenotype in a *ΔGr28* deletion mutant lacking both Gr28a and Gr28b (***Figure 4 – figure supplement 1***).

To confirm Gr28b is necessary for responses to CO_2_ in td neurons, we used a more selective Gr28b null mutant line, *Gr28b^ΔSL06^*, generated by scarless genome engineering (Feng et al., 2021). Loss of Gr28b was confirmed by Southern blot analysis (Feng et al., 2021). The mutant line was homozygous viable and showed no overt phenotypes by third instar larval stage. We used 2 minutes 20% CO_2_ or 30 seconds 79-80% CO_2_ as the stimuli while holding O_2_ constant at 20-21%. We found that A3 v’td2 responses to CO_2_ were significantly reduced in *Gr28b^ΔSL06^* mutants in both conditions (***Figure 4B-D***). Notably, responses were also reduced in *Gr28b^ΔSL06^* heterozygous mutants compared to control larvae (***Figure 4B-C***), suggesting dosage sensitivity in the response. We also tested whether Gr28b is involved in oxygen sensing by exposing *Gr28b^ΔSL06^* mutants to 1 minute 0% O_2_. We found that A1 v’td2 neurons showed normal responses to anoxia in *Gr28b^ΔSL06^*mutant animals (***Figure 4E***). Altogether, our results are consistent with a requirement for Gr28b in CO_2_-induced responses, but not in O_2_ responses, in td neurons.

### Rescuing Gr28b.c expression restores responses to CO_2_ in td neurons in Gr28b^ΔSL06^ mutants

The Gr28b locus is complex and encodes five isoforms (a, b, c, d, e), each of which has a unique transcriptional start site and exon, spliced with two common exons (Montell, 2013). Both of the Gr28b RNAi lines that we used target the common exons, and the *Gr28b^ΔSL06^* mutant deletes all five isoforms. We examined the expression pattern of all five Gr28b isoforms in larva (Thorne & Amrein, 2008). Of the five isoforms, we only found the *Gr28b.c-Gal4* reporter labels td neurons, suggesting that Gr28b.c may mediate the response in CO_2_-sensitive td neurons (***Figure 3 – figure supplement 1***). To test the functions of Gr28b.c specifically and examine the potential functions of other Gr28b isoforms, we rescued the expression of each of the five isoforms in *Gr28b^ΔSL06^*mutant animals. We found that rescue of Gr28b.c strongly restored responses to 79-80% CO_2_ in *Gr28b^ΔSL06^* mutant animals (***Figure 5A***). Restoration of Gr28b.e moderately increased responses to CO_2_ above *Gr28b^ΔSL06^* mutant animals but not to the same extent as Gr28b.c rescue (***Figure 5A***).

**Figure 5:**
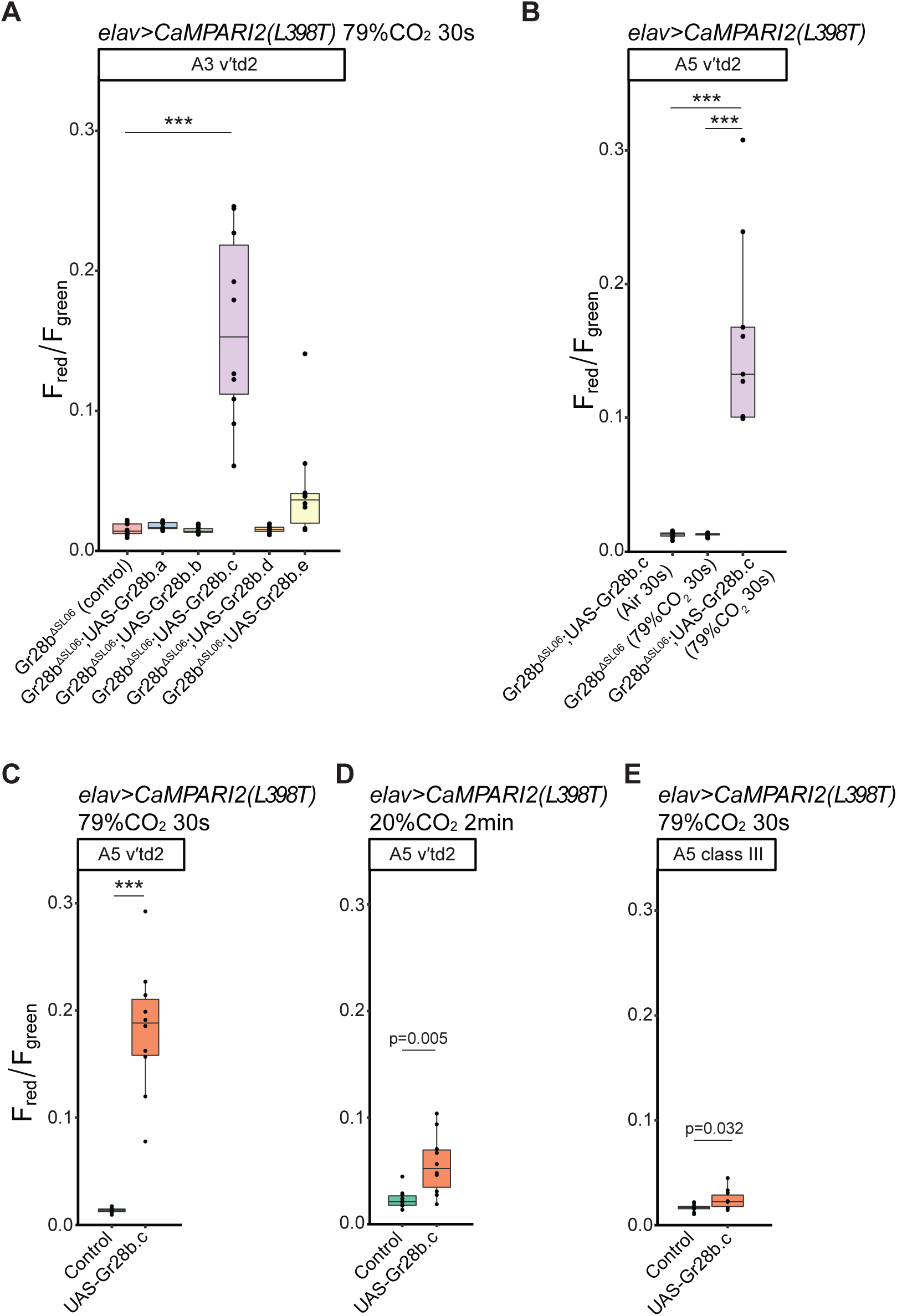
Ectopic Gr28b.c expression rescues responses to CO2 in td neurons. **A.** Quantification of red/green CaMPARI2 fluorescence in A3 CO2-sensitive v’td2 neurons in *Gr28b^ΔSL06^*animals (control) and ectopic expression of five Gr28b isoforms under the control of *elav-Gal4* in *Gr28b^ΔSL06^* background in response to 30s of 79% CO2. Ectopic Gr28b.c expression significantly rescued the responses of A3 v’td2 neurons to high CO2 compared to *Gr28b^ΔSL06^*. Ectopic Gr28b.e expression moderately increased the response of A3 v’td2 to high CO2 compared to *Gr28b^ΔSL06^*. n=10-11 animals per condition. **B.** Quantification of red/green CaMPARI2 fluorescence in A5 v’td2 neurons in *Gr28b^ΔSL06^*, and *Gr28b^ΔSL06^* with ectopic Gr28b.c expression animals. *Gr28b^ΔSL06^* with ectopic Gr28b.c expression larvae exposed to 30s air, and *Gr28b^ΔSL06^* larvae exposed to 30s 79% CO2 (same animals as in **A**) showed no td neuron activity. *Gr28b^ΔSL06^*with ectopic Gr28b.c expression larvae exposed to 30s 79% CO2 (same animals as in **A**) showed significantly increased activities in all v’td2 neurons compared to the other two conditions. n=10-11 animals per condition. **C-D.** Quantification of red/green CaMPARI2 fluorescence in A5 v’td2 neurons in control and upon Gr28b.c overexpression in response to 30 seconds 79% CO2 and 2 minutes 20% CO2. Gr28b.c overexpression resulted in significant increases in fluorescence ratio in v’td2 neurons in high CO2. n=10 animals per condition. **E.** Quantification of red/green CaMPARI2 fluorescence in the A5 class III somatosensory neuron in control and misexpression conditions in response to 30 seconds of 79% CO2. Ectopic expression of Gr28b.c did not induce response to CO2. n=10 animals per condition.

Because rescuing Gr28b.c expression restored responses to CO_2_ in a *Gr28b^ΔSL06^*mutant background we sought to test if Gr28b.c was sufficient to confer responses to different levels of external CO_2_ in normally non-responsive cells (***Figure 5B***). As described above, A4-A7 v’td2 neurons are labelled by *Gr28b.c-Gal4* but did not show as consistent responses to CO_2_ using the CaMPARI2 reporter. Focusing on A5 v’td2, we found that overexpression of Gr28b.c in control or *Gr28b^ΔSL06^*mutant backgrounds induced strong responses to CO_2_ (***Figure 5B-D***). Moreover, ectopic expression of Gr28b.c induced strong responses in *Gr28b.c-Gal4* non-expressing v’td1 neurons after 30 seconds of 79-80% CO_2_ exposure (***Figure 5 – figure supplement 1A-B***). However, other peripheral somatosensory neurons (class III da neuron) do not acquire as strong responses to CO_2_ upon ectopic Gr28b.c expression, suggesting the induced responses are specific to td neurons (***Figure 5E***). Our results suggest that high levels of Gr28b.c expression are sufficient to confer CO_2_ sensitivity to td neurons.

### Gr28a and Gr33a do not mediate responses to CO_2_ in td neurons

Our Gr28b rescue experiments strongly suggested that Gr28b.c participates in responses to increased CO_2_ levels in td neurons. We note, however, that *Gr28b.c-Gal4* is expressed by all v’td2 neurons but only a subset is responsive to increased CO_2_ levels. We next asked why other *Gr28b.c-Gal4*-positive td neurons did not normally respond to increased CO_2_ levels. One possibility is that *Gr28b.c-Gal4* expression does not reflect expression of *Gr28b.c* in all td neurons. We also considered the possibility that other gustatory receptors influence Gr28b.c activity. The Gr28b.c reporter is co-expressed with reporters for Gr28a and Gr89a only in neurons that show CO_2_ sensitivity (Qian et al., 2018). We wondered whether the expression of these other two GRs may be necessary for Gr28b.c-mediated responses to CO_2_. To test this, we ectopically expressed Gr28a pan-neuronally, such that *Gr28b.c-Gal4* positive td neurons now co-express Gr28a, Gr28b.c, and Gr89a (***Figure 6A***). Using A5 v’td2 neuron as a representative, we found that td neurons that do not normally respond strongly to CO_2_ also did not acquire sensitivity to CO_2_ with ectopic expression of Gr28a, suggesting that additional factors beyond Gr28a and Gr89a are involved (***Figure 6B***).

**Figure 6:**
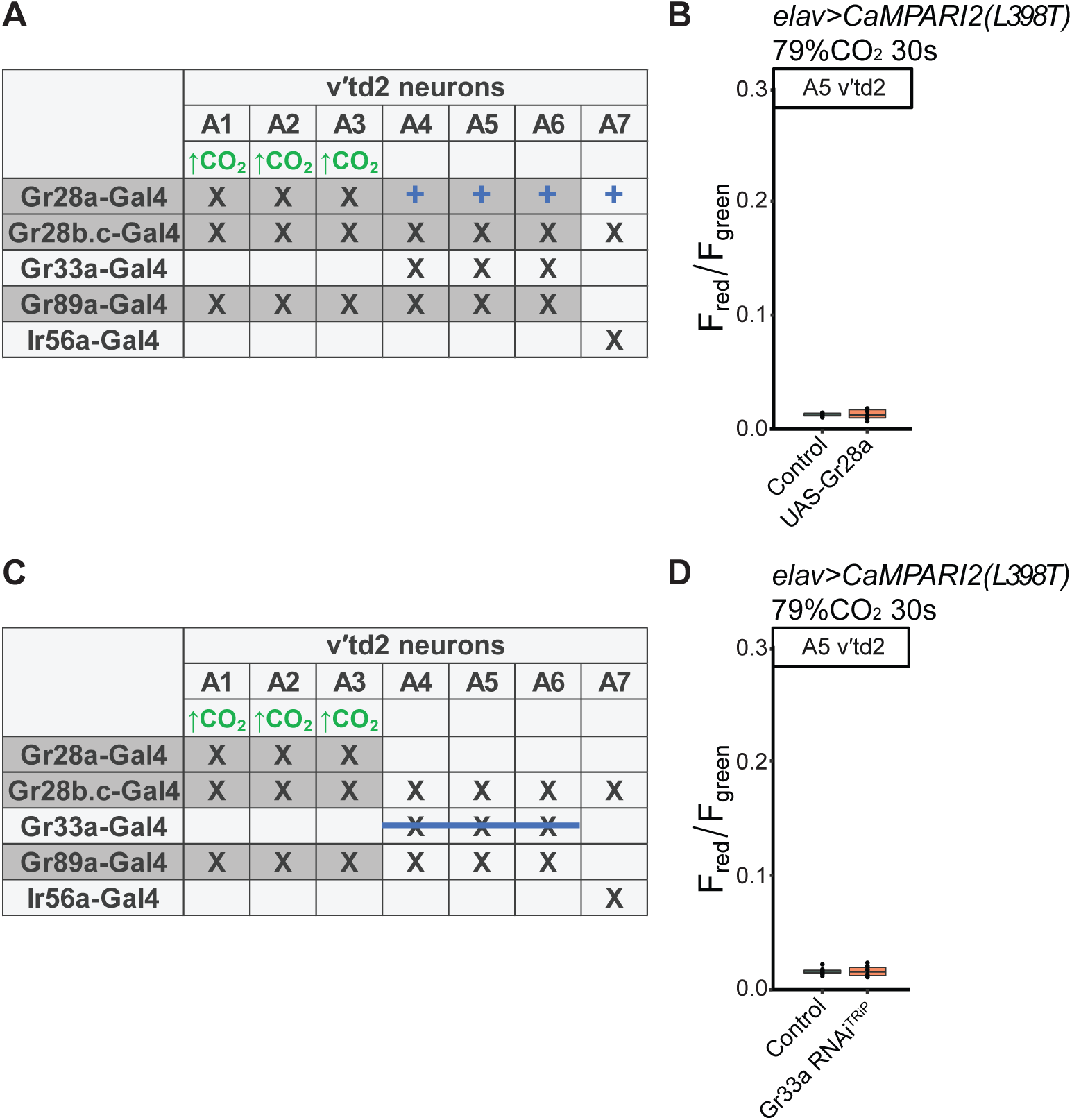
Gr28a and Gr33a do not mediate responses to CO2 in td neurons. **A.** Summary of experimental manipulation of chemosensory receptor expression in v’td2 neurons in panel **B**. Blue plus signs indicate ectopic expression of Gr28a. Gray boxes indicate co-expression of Gr28a, Gr28b.c, and Gr89a. **B.** Quantification of red/green CaMPARI2 fluorescence in v’td2 neurons in control and Gr28a misexpression animals in response to 30s 79% CO2. A5 v’td2 neurons in Gr28a misexpression animals showed no detectable increase in response to high CO2 level. n=10-11 animals per condition. **C.** Summary of experimental manipulation of chemosensory receptor expression in v’td2 neurons in panel **D**. Blue strikethrough indicates knockdown of Gr33a. **D.** Quantification of red/green CaMPARI2 fluorescence in v’td2 neurons in control and Gr33a knockdown animals in response to 30s 79% CO2. A5 v’td2 neurons showed no detectable increase in fluorescence in response to high CO2 when Gr33a was knocked down using RNAi. n=10 animals per condition.

GR-GR inhibitory interactions have been observed in bitter gustatory receptor neurons in *Drosophila* (Dweck & Carlson, 2020). Thus, we reasoned that Gr33a could interfere with the effect of other receptors (Delventhal & Carlson, 2016; Sung et al., 2017). We knocked down the expression of Gr33a using RNAi and examined responses to CO_2_ in the A5 v’td2 neuron (***Figure 6C***). We observed no difference in responses to 79-80% CO_2_ between control animals and Gr33a knockdown animals. With the caveat that knock down could be incomplete, these data suggest that inhibitory interactions between Gr33a and the other GRs do not prevent responses to CO_2_ (***Figure 6D***). Mechanisms other than co-expression of specific other GRs, or inhibition by a GR expressed in a complementary fashion therefore likely account for the observed selectivity of CO_2_ sensitivity.

### td neurons are activated by prolonged submersion

Given the responses of td neurons to heightened CO_2_ levels we hypothesized that a subset of td neurons helps to monitor the internal gas concentration under natural conditions (e.g. feeding). We therefore assessed whether td neurons are activated when larvae burrow for prolonged periods. Previous studies have shown that when oxygen is completely removed, *Drosophila* larvae survive and move for at least the first 40 minutes (Callier et al., 2015). To simulate natural feeding conditions, we used 0.8% agar gel, which falls within the range of substrate stiffness encountered by feeding larvae (Kim et al., 2017). When larvae are completely submerged, their spiracles are closed, preventing gas exchange (Manning & Krasnow, 1993). To capture neural activity following prolonged submersion, we placed larvae in agar plates to prevent gas exchange for 5 or 10 minutes. Plates were sealed with parafilm to prevent larvae from exiting the agar. Immediately after the submersion period, we applied UV for 30 seconds to enable CaMPARI2 photoconversion, when the animals are still sealed in agar gel (***Figure 7A***). As a control, we applied 30 seconds of UV to larvae right after they completely burrow into an agar plate covered in parafilm. After 5 minutes and 10 minutes of submersion, A1-A3 td neurons showed significantly higher responses than controls (***Figure 7B-C*)**. v’td2 neurons in A4 and below showed modest, but significant, responses.

**Figure 7.**
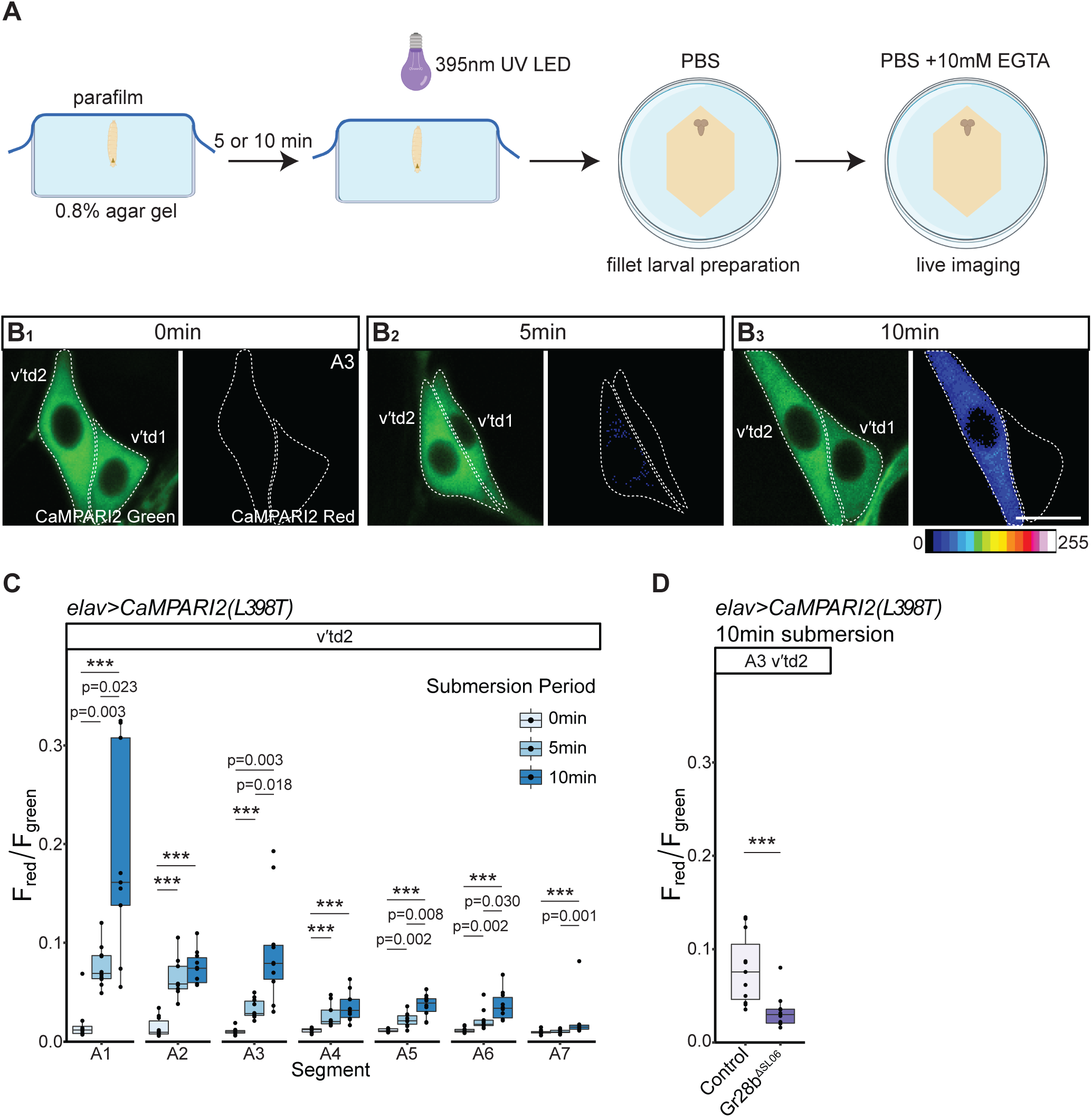
td neurons are activated by prolonged submersion. **A.** Schematic of protocol for post-hoc neural imaging of responses to prolonged submersion. (Created with BioRender.com). **B.** Green and red CaMPARI2 fluorescence in A3 td neurons in response to prolonged submersion. (Red channel shown in pseudo color). Dorsal is top, and anterior is to the left. **B1.** Larvae exposed to UV immediately after submersion showed little to no green to red CaMPARI2 photoconversion in td neurons. **B2-3.** Larvae submerged for 5 or 10min then exposed to UV showed increased green to red photoconversion in A3 v’td2 neuron but not in v’td1 neuron. **C.** Quantification of red/green CaMPARI2 fluorescence in all v’td2 neurons across segments A1-A7 in response to prolonged submersion. The three gas-sensitive td neurons (A1-A3 v’td2) responded to 5 or 10min of submersion under agar gel. Other slow responding v’td2 showed modest, but significant, responses to submersion. n=8-10 animals per condition. **D.** Quantification of red/green CaMPARI2 fluorescence in A3 CO2-sensitive v’td2 in response to prolonged submersion in control and *Gr28b^ΔSL06^* animals. *Gr28b^ΔSL06^*larvae showed significant decrease in response to prolonged submersion compared to control animals. n=10-11 animals per condition.

Since Gr28b.c mediates responses to environmental CO_2_, we next asked whether it also mediates responses to internally generated CO_2_ build-up in submerged larvae. We focused on A3 v’td2 neurons since it only responds to CO_2_-related changes. We submerged *Gr28b^ΔSL06^* mutant animals in 0.8% agar gel for 10 minutes and monitored the responses in A3 v’td2 neurons using CaMPARI2 as described above. We found that *Gr28b^ΔSL06^* mutants showed significantly reduced responses to prolonged submersion compared to control larvae (***Figure 7D***). Altogether, these results suggest that td neurons are activated when gas exchange becomes impaired in natural-like conditions, and that Gr28b mediates these responses.

## DISCUSSION

The diverse sensory functions of interoceptive systems are relatively less explored than systems that primarily monitor external stimuli. Here, we explored the morphology and functions of internal td neurons in *Drosophila* and found that partially overlapping subsets of td neurons respond to a decrease in O_2_ and increase in CO_2_ levels. We identified a role for the guanylyl cyclases, Gyc88E and Gyc89Db, in mediating td responses to decreased O_2_. We also focused on the role of Gr28b in CO_2_ sensing. We provide genetic evidence that Gr28b mediates the response of td neurons to CO_2_ and that a specific isoform, Gr28b.c, can endow td neurons with sensitivity to high CO_2_ levels. We also provide evidence that these gas-sensitive neurons are activated by prolonged submersion in agar gel, demonstrating a natural-like condition in which td neurons are activated.

Our morphological characterization of td dendrites suggests that they are internal to the animal’s body and not directly exposed to the tracheal lumen. Electron microscopy along the whole td dendrite and tracheal tube would be needed to have a full description of the dendrite-tracheal association. This could be accomplished using the recently-described full larval SEM volume (Richter et al., 2025; Schoofs et al., 2024). Given the close association and signalling between dendrites of body wall sensory neurons and epidermis, it is also notable that td dendrites have a clear substrate preference for tracheal epithelial cells (Han et al., 2012; Kim et al., 2012). The molecules that underlie this dendrite substrate preference will be an interesting topic for future investigations since our results suggest that this association is important for td sensory functions.

Gas-sensing in *Drosophila* had previously been studied in sensory neurons with external sensilla (Jones et al., 2007; Kwon et al., 2007). By contrast, td neuron dendrites are internal and exposed to the hemolymph along their length. In *Drosophila* larvae, a pair of chemosensory neurons in the head terminal organ express Gr21a and Gr63a which together function as CO_2_ receptors. These CO_2_ receptors are very sensitive to small changes in gas concentrations, allowing animals to respond rapidly to avoid a potential harmful environment. Ablation of these neurons abolish behavioral avoidance to 1% CO_2_ (Faucher et al., 2006). In addition, a population of gustatory neurons in the adult responds to aqueous and gas phase (10%-100%) CO_2_ (Fischler et al., 2007). Unlike the olfactory neurons, these gustatory neurons lack Gr21a/63a expression and detect CO_2_ through Ir25a/56d/76b (Sanchez-Alcaniz et al., 2018). Here we describe another CO_2_ sensing system comprising td neurons that are activated by high levels of CO_2_ in the environment (10-100%) and provide evidence that Gr28b mediates these responses.

During respiration, O_2_ is constantly consumed and CO_2_ produced. In the respiratory (tracheal) system of *Drosophila* larvae, gas exchange occurs through passive diffusion (Manning & Krasnow, 1993). The high external CO_2_ concentrations used in our experiments may disrupt the normal CO_2_ gradient across the spiracles, impairing gas exchange and causing CO_2_ to accumulate internally. Therefore, the increases in tracheal CO_2_ may be much lower than the CO_2_ concentration we applied externally. td neurons are delicately positioned so they may detect physiological changes in tracheal epithelial cells or in the hemolymph near tracheal branches, which is vastly different from the environment that external CO_2_-sensing neurons are exposed to, perhaps necessitating a different molecular mechanism for detecting CO_2_ than the well-established Gr21a and Gr63a.

A recent study has linked td neurons to corazonin and diuretic hormone producing neuroendocrine cells (Huckesfeld et al., 2021). By monitoring internal chemical states, td neurons could conceivably promote stress responses and excretion of acidified hemolymph to maintain homeostasis. A developmental study showed that A1-A3 td neurons labelled by Gr28a are necessary for the progression of larval growth (Ohhara & Yamanaka, 2022), indicating another possible scenario where td neurons are required for monitoring developmental status of the larva. Given the activation of td neurons during prolonged submersion, one trigger for td neuron activity could be respiratory stress that occurs during feeding. Larvae show cooperative feeding behavior that involves repeated diving into food and surfacing. By creating feeding cavities deep below the food surface, cooperative feeding brings larvae closer to the most nutritious food (Dombrovski et al., 2017). However, during these dives, larvae forgo respiration. Transitioning between feeding and surfacing modes balances energy expenditure and nutrient acquisition, while reducing the risk of desiccation and respiratory distress (Dombrovski et al., 2017; Kim et al., 2017). Notably, digging and diving behavior is suppressed when posterior spiracles are obstructed, raising the possibility that diving-induced hypoxia or hypercapnia may also promote surfacing (Kim et al., 2017). We speculate that during cooperative feeding the cavity microenvironment could be depleted in O_2_ and enriched in CO_2_ due to larval respiration, making these relatively stressful microenvironments that require attention to internal state. It will be important to test roles for td neurons in these feeding behaviors and responses to respiratory stresses.

The Gr28b locus generates five isoforms, Gr28b.a-Gr28b.e. The functions of most Gr28b isoforms - including Gr28b.c that is expressed in td neurons - are not known. Gr28b proteins appear to have non-canonical roles as gustatory receptors. For example, the Gr28b.d isoform functions as a warmth sensor (Ni et al., 2013), and an unidentified Gr28b isoform is required for light-avoidance in nociceptive (class IV da) sensory neurons (Xiang et al., 2010). We observed that *Gr28b.c-Gal4* labels class IV da neurons, suggesting Gr28b.c may be involved in the noxious light transduction in nociceptive neurons. However, the study also pointed out that the light-avoidance response requires TrpA1 together with Gr28b.c (Xiang et al., 2010), suggesting Gr28b.c is involved in, but not sufficient for, noxious light-transduction. The way that Gr28b.c is involved in the noxious light-transduction is still not understood. In addition, another study in *Drosophila* adults showed that Gr28b.c is required for the avoidance of a bitter compound, saponin, suggesting a common theme for Gr28b.c in detecting noxious stimuli. Our results suggest that Gr28b.c is necessary for internal CO_2_-detection, and ectopic Gr28b.c expression strongly increases responses to CO_2_ in td neurons, but very weakly in other sensory neurons (e.g. class III da neurons) residing on the body wall. One possibility is that the location of td sensory processes on the trachea impact responsiveness. Alternatively, or in addition, other factors present in td neurons may work in combination with Gr28b to impart responsiveness. Carbon dioxide in water can undergo interconversion to form bicarbonate and a free proton catalysed by carbonic anhydrases (Guyenet & Bayliss, 2015). Thus, CO_2_ may be detected inside the body as increases in H^+^ (i.e. low pH) or HCO^−^_3_ concentration rather than directly as CO_2_. The exact ligands and mechanisms of sensing will be important questions for future studies.

Out of the three td neurons that responded strongly to increases in CO_2_, two neurons also responded to decreases in O_2_. The two O_2_-sensing td neurons express molecular oxygen sensors soluble guanylyl cyclases Gyc88E and Gyc89Db, and our data suggest that these two soluble guanylyl cyclases mediate td responses to hypoxia. Gyc88E can form homodimers, or form heterodimers with either Gyc89Da or Gyc89Db. By contrast, Gyc89Da or Gyc89Db require Gyc88E for their activity (Morton, 2004). We observed that ectopically co-expressing Gyc88E and Gyc89Db induced higher hypoxia responses in td neurons compared to ectopically expressing Gyc88E alone. One interpretation of these results is that Gyc88E/Gyc89Db co-expression permits formation of heterodimers that are more sensitive to hypoxia compared to Gyc88E alone (Morton, 2004). Since these tracheal neurons are situated internally, they could be well-positioned to detect and elicit responses to dangerously low internal O_2_ levels during prolonged diving into food (Kim et al., 2017), as compared to tail sensory neurons which could be more affected by ambient O_2_ levels.

Our imaging and anatomical results together show that CO_2_- and O_2_-sensitive td neurons project axons to the SEZ (Qian et al., 2018). Previous studies showed that olfactory CO_2_-sensing neurons in the larva target the larval antennal lobe (Python & Stocker, 2002). Thus, internal and external CO_2_-sensitive neurons may have different downstream effects. Given that both td neuron axons and gustatory sensory axons target the SEZ, internal chemosensation and gustation may have similar downstream effects. Future studies that manipulate td neuron activity or that of their downstream CNS targets should provide insights into the behavioral or physiological functions of td gas-sensing circuits.

## ACKNOWLEDGMENTS

We thank Drs. Benjamien Moeyaert, Seok Jun Moon, Hubert Amrein, John Carlson, Paul Garrity, David Morton and the Bloomington Stock Center for fly stocks, Dr. Siqian Feng for advice on scarless genome engineering, Drs. John Carlson and Hany Dweck for comments on the manuscript, and members of the Grueber lab and Drs. Mimi Shirasu-Hiza, Richard Mann and Laura Duvall for advice and support. Research reported in this publication was supported by the National Institute of Neurological Disorders and Stroke of the National Institutes of Health under Award Number R21NS105507 and 5R01NS061908 to WBG. The content is solely the responsibility of the authors and does not necessarily represent the official views of the National Institutes of Health.

## MATERIALS AND METHODS

### Drosophila stocks

Animals were reared using standard methods. Animals of either sex were analysed at the late third instar larval stage. The following stocks were used, and were obtained from Bloomington Drosophila Stock Center and Vienna Drosophila Resource Center unless otherwise indicated: *w^1118^* (VDRC isogenic host strain; VDRC ID 60000), *Gyc88E-Gal4* (*490-Gal4*; RRID:BDSC_63353), *Gyc89Da-Gal4*; *Gyc89Db-Gal4* (gifts from David B. Morton, Oregon Health & Science University), *nompc-Gal4* (RRID:BDSC_36361), *piezo-Gal4* (RRID:BDSC_58771), *breathless-Gal4* (RRID:BDSC_78328), *Gr28a-Gal4* (RRID:BDSC_57613), *Gr28a-QF2^G4H^* (generated using the HACK method) (Lin, 2016}, *Gr28b.a-Gal4* (RRID:BDSC_57615; RRID:BDSC_58369), *Gr28b.b-Gal4* (RRID:BDSC_57616; RRID:BDSC_57617), *Gr28b.c-Gal4* (RRID:BDSC_57618; RRID:BDSC_57619), *Gr28b.d-Gal4* (RRID:BDSC_57620), *Gr28b.e-Gal4* (RRID:BDSC_57621), *Gr33a-Gal4* (RRID:BDSC_57623; RRID:BDSC_57624), *Gr89a-Gal4* (RRID:BDSC_57676), *Ir25a-Gal4* (RRID:BDSC_41728), *Ir56a-Gal4* (RRID:BDSC_60704), *Ir76b-Gal4* (RRID:BDSC_41730), *elav-Gal4* (RRID:BDSC_8760; RRID:BDSC_8765), *GMR35B01-Gal4* (RRID:BDSC_49898), *UAS-mCD8-GFP* (RRID:BDSC_5137; RRID:BDSC_32194), *10xQUAS-6xGFP, 5xUAS-mtdt-3HA* (Lin and Potter, 2016; Potter et al., 2010), *UAS-CaMPARI2(L398T)* (gift from Benjamien Moeyaert, Schreiter lab, Janelia Research campus), *Gr28a^1^* (gift from Seok Jun Moon, Yonsei University), *Gyc88E GD11230* (RRID:FlyBase_FBst0454241), *Gyc88E TRiP.HM05096* (RRID:BDSC_28608), *Gyc89Db TRiP.HM05207* (RRID:BDSC_29529), *Gr28b TRiP.HMJ30111* (RRID:BDSC_63545), *Gr28b KK105653* (RRID:FlyBase_FBst0473600), *ΔGr28 30i* (gift from Hubert Amrein, Texas A&M University), *Gr33a TRiP.HMJ30017* (RRID:BDSC_62940), *Gr89a TRiP.HMS05359* (RRID:BDSC_64023), *UAS-Gyc88E; UAS-Gyc89Db* (gifts from David B. Morton, Oregon Health & Science University), *UAS-Gr28a* (gift from John Carlson, Yale University), *UAS-Gr28b.a*; *UAS-Gr28b.b*; *UAS-Gr28b.c*; *UAS-Gr28b.d*; *UAS-Gr28b.e* (gifts from Paul Garrity, Brandeis University).

For CaMPARI2 knock-down or overexpression experiments, we used the progeny of *elav-Gal4, UAS-CaMPARI2(L398T)* flies crossed with *w^1118^*flies as control. For experimental groups, we crossed *elav-Gal4, UAS-CaMPARI2(L398T)* flies to *UAS-RNAi, UAS-Gyc* or *UAS-Gr* fly lines and examined CaMPARI2 responses in the progeny.

### Generation of *Gr28b* mutant fly lines

*Gr28b^ΔSL06^* mutant fly lines were generated using a scarless engineering method for the *Drosophila* genome as previously described (Feng et al., 2021).

### Immunohistochemistry

Immunohistochemistry was performed as previously described (Matthews et al., 2007). Third instar larvae were dissected in 1X PBS, fixed in 4% paraformaldehyde (Electron Microscopy Sciences; CAS #30525-89-4) in PBS for 15 minutes, rinsed three times in PBS-TX (0.3% Triton X-100 in PBS), and blocked for 1 hour at 4°C in normal donkey serum (1:20; Jackson Immunoresearch; RRID: AB_2337258). Primary antibodies used were chicken anti-GFP (1:1000; Abcam; RRID: AB_300798), rabbit anti-DsRed (1:500; Takara Bio, RRID:AB_10013483), goat anti-HRP (1:200; Jackson Immunoresearch; RRID: AB_2338952), mouse anti-Coracle (1:10; Developmental Studies Hybridoma Bank; RRID:AB_1161642, RRID:AB_1161644) and mouse anti-Elav (1:10; Developmental Studies Hybridoma Bank; RRID:AB_528217). Secondary antibodies used were Alexa Fluor 488 donkey anti-chicken (1:200; Jackson Immunoresearch; RRID: AB_2340375), Rhodamine Red-X donkey anti-rabbit (1:200; Jackson Immunoresearch; RRID: AB_2340613), Alexa Fluor 647 donkey anti-goat (1:200; Jackson Immunoresearch; RRID: AB_2340437), Rhodamine Red-X donkey anti-goat (1:200; Jackson Immunoresearch; RRID: AB_2340423), Rhodamine Red-X donkey anti-mouse (1:200; Jackson Immunoresearch; RRID: AB_ AB_2340831), and Alexa Fluor 647 donkey anti-mouse (1:200; Jackson Immunoresearch; RRID: AB_2340862).

Immunohistochemistry for anti-HRP without detergent was performed as previously described (Kim et al., 2012). Fixed animals were rinsed three times with PBS and labeled with goat-anti-HRP (1:200) in PBS for 30 minutes at room temperature. The animals were rinsed and washed in PBS for 30min, then in PBS-TX for 30min, followed by regular immunohistochemistry with the other primary and secondary antibodies.

### CaMPARI2 imaging

For CaMPARI2 imaging of neuronal responses to gaseous stimuli, individual late third-instar larvae were washed in PBS buffer, dried on a Kimwipe, and transferred to a detached centrifuge tube lid. The larva was covered with a petri dish lid and placed over a spacer ring and a FlyPad (Genesee Scientific). A 395nm UV LED (LEDwholesalers, 7202UV395) was positioned on top of the spacer ring. The UV LED was used at 3.5V, supplied by a bench power supply (TENMA, 72-7245). Gas mixtures (Tech Air; Airgas) were delivered to the FlyPad at 5L/min through a benchtop Flowbuddy (Genesee Scientific). Following coincident application of gaseous stimuli and UV light for the specified duration, the larva was immediately immersed in PBS and dissected in a fillet larval preparation. The buffer was replaced with 2mL of fresh PBS with 10mM EGTA (pH 7.3-7.5) (CAS #67-42-5) and the larva was immediately imaged using a Zeiss LSM 700 with a 40x water-immersion Plan-Apochromat objective lens (N.A.=1.0). Green CaMPARI2 fluorescence was imaged using the both the 488nm and low power 405nm (at 5% power for improved signal, see (Fosque et al., 2015)) laser lines. Similarly, the red CaMPARI2 fluorescence was imaged using both the 555 nm and low power 405 nm laser lines (Fosque et al., 2015). Single 5μm thick sections were acquired. One pair of td neurons per segment was acquired for each animal.

For CaMPARI2 imaging of neuronal responses to prolonged submersion, individual late third-instar larvae were placed in 35 x 10 mm petri dishes (Falcon) filled completely with 0.8% agar gel. The agar gel was gently stabbed to allow larvae to burrow in. After larvae completely burrowed into the gel, the agar plate was tightly sealed with parafilm with all air bubbles carefully squeezed out with a fine brush. A 395nm UV LED was applied from above for 30 sec either starting immediately or after the specified submersion duration. After CaMPARI2 photoconversion, larvae were immediately removed from the agar gel, dissected in PBS and imaged in PBS with 10mM EGTA, as described above.

### CaMPARI2 image analysis

Images were acquired for one pair of td neurons in each segment along the length of the larva. All images were analysed blind to genotype or stimulus. An ROI was manually drawn around the cell body of each td neuron, excluding regions obscured by nearby tissues, in Fiji (Schindelin et al., 2012). The mean green fluorescence (F_green_) and red fluorescence (F_red_) were measured for the selected ROI. The same ROI was copied and moved to a region next to td neuron and not overlapping with nearby tissues. For background subtraction, the mean green fluorescence (F_background green_) and red fluorescence (F_background red_) were measured for the new ROI. The CaMPARI ratio was calculated as (F_red_ - F_background red_) / (F_green_ - F_background green_) (Edwards et al., 2020).

### Statistical Analyses

Statistical analyses were conducted using RStudio Version 1.2.5033. Box plots show first to third quartiles and median as middle line. The whiskers indicate 1.5 inter-quartile range. Shapiro-Wilk normality test was used to determine normal distribution for all experiments. Depending on the result, either parametric or non-parametric tests were selected. For comparing two groups, Welch’s t-test or Mann-Whitney U test was selected for quantifications of CaMPARI2 fluorescence ratio and test for statistical significance between control and experimental conditions. For comparing three or more groups, Welch’s ANOVA test followed by Games-Howell test or paired Welch’s t-test, or Kruskal-Wallis test followed by pairwise Wilcoxon rank sum test was selected for testing statistical significance between control and experimental conditions. Bonferroni correction was performed to obtain adjusted p values. All comparisons made were within a segment. p values < 0.05 are indicated in the figures, and p values < 0.001 are represented by ***. See figure legends for sample sizes.

---

**Figure 1 – figure supplement 1:**
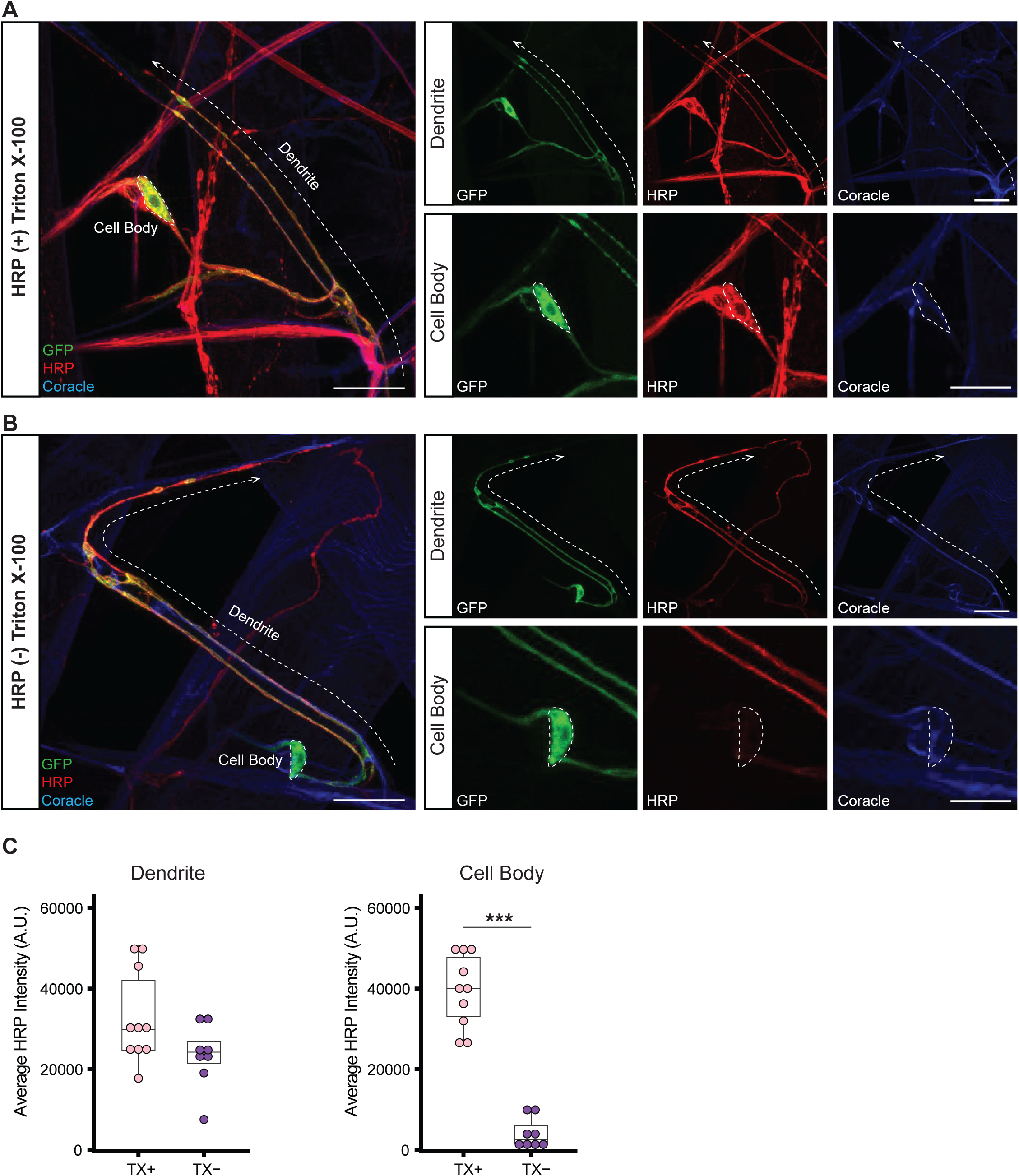
Additional characterization of td dendrite association with tracheae. **A.** td neuron cell bodies and dendrites labeled with GFP (green), HRP (red), and Coracle (blue), all after treatment with Triton X-100. Cell bodies (dashed outline), dendrites (dashed line), and tracheae are visible in insets. **B.** Immunolabeling of HRP (red channel) prior to Triton X-100 permeabilization step identifies membrane protected cell bodies but no membrane protection of dendrites. **C.** Average HRP labeling along dendrites is unaffected by Triton X-100 treatment. Cell body HRP labeling is significantly lower without Triton X-100 permeabilization.

**Figure 2 – figure supplement 1:**
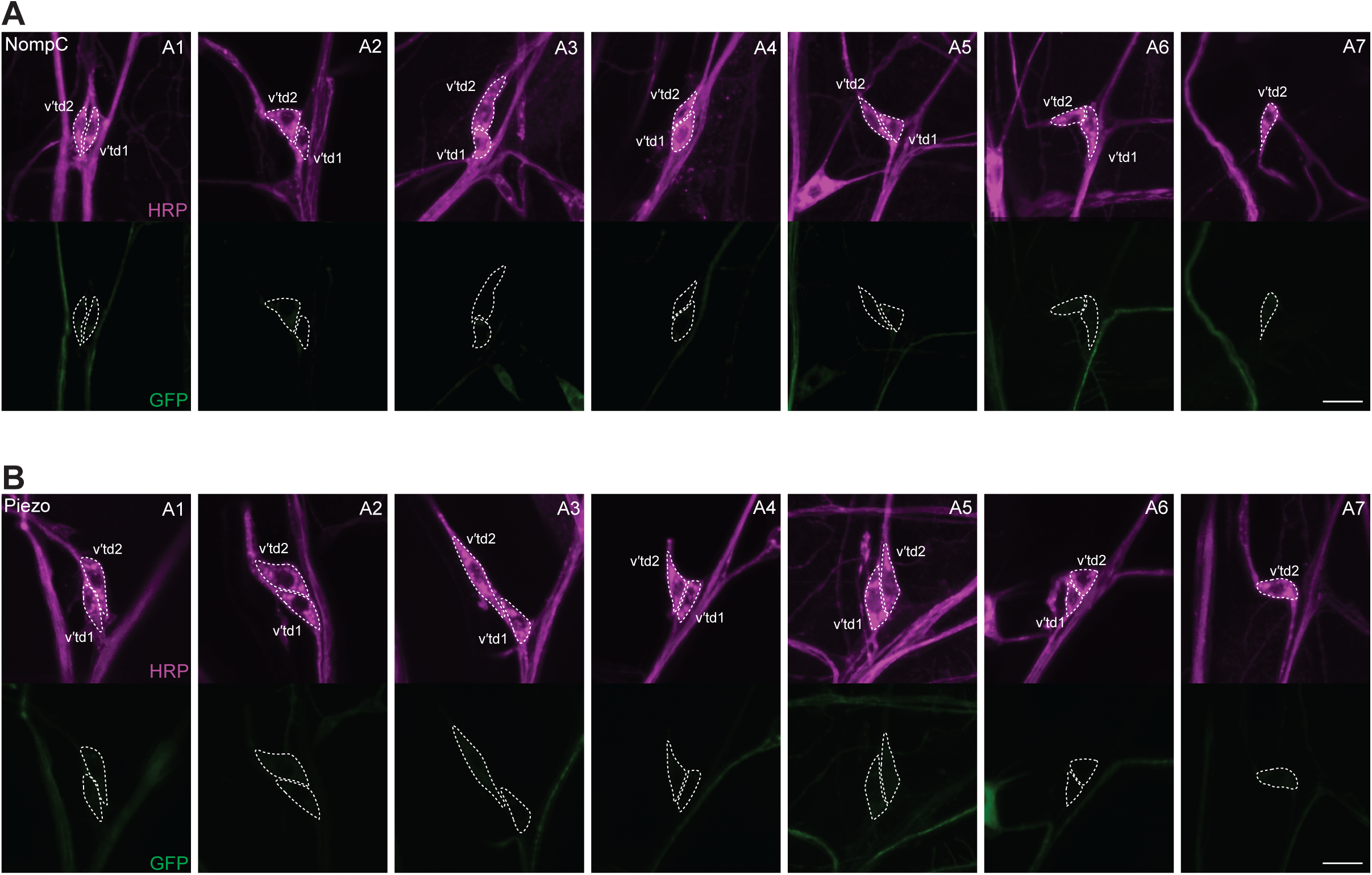
Mechanosensory receptor reporters are not detected in td neurons. Images of A1-A7 td neurons in **(A)** *NompC-Gal4>UAS-mCD8::GFP* and **(B)** *Piezo-Gal4>UAS-mCD8::GFP* larvae. Anti-GFP is in green and anti-HRP in magenta. No td neurons showed detectable expression of either reporter line. Dorsal is top, and anterior is to the left in all images. Scale Bar, 10μm for all images.

**Figure 2 – figure supplement 2:**
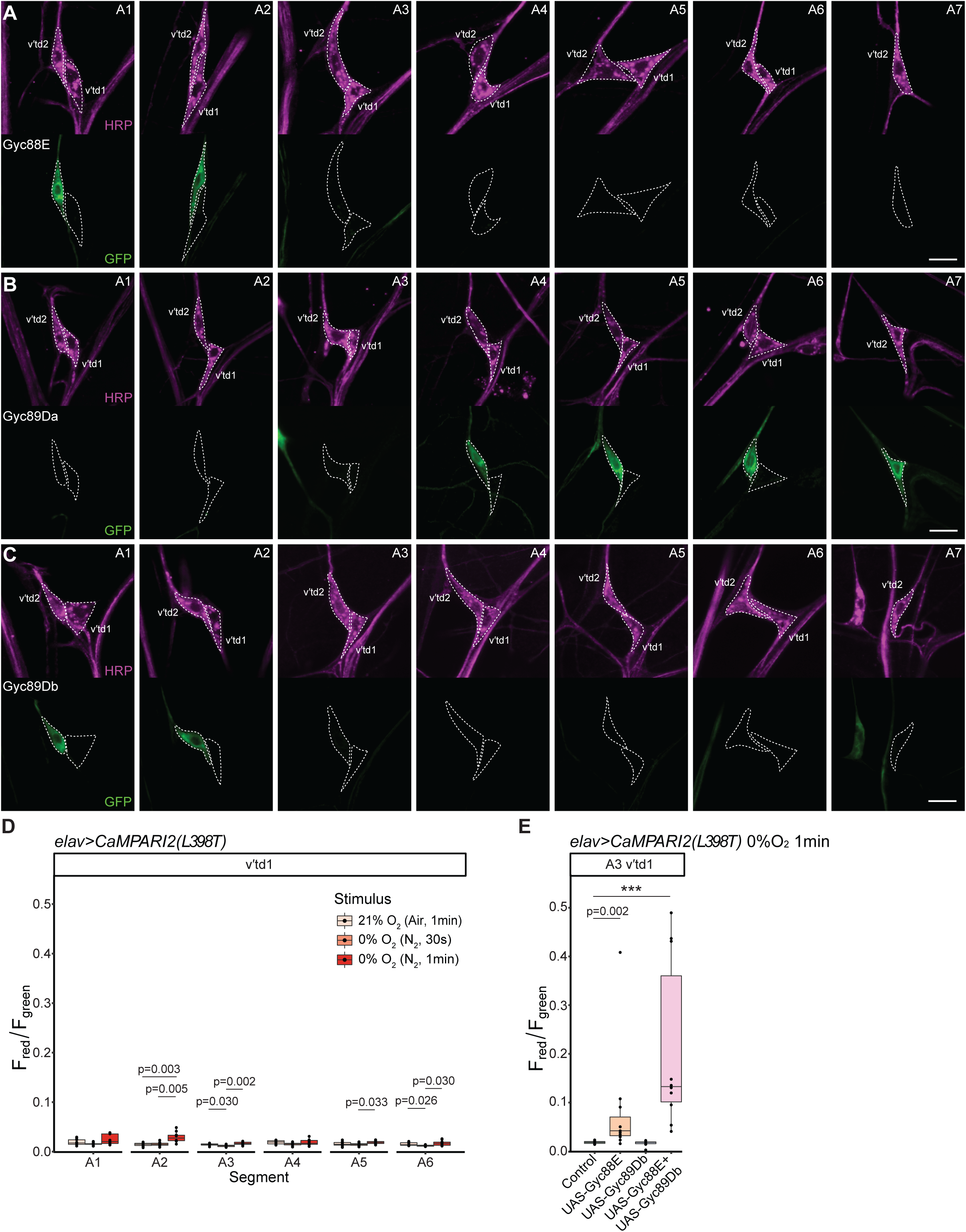
Expression of Gyc-Gal4 lines in td neurons. Images of A1-A7 td neurons in **A.** *Gyc88E-Gal4>UAS-mCD8::GFP*, **B.** *Gyc89Da-Gal4>UAS-mCD8::GFP*, and **C.** *Gyc89Db-Gal4>UAS-mCD8::GFP* larvae. Anti-GFP is shown in green and anti-HRP in magenta. **D.** v’td1 neurons are insensitive to anoxic conditions. **E.** Misexpression of *UAS-Gyc88E* alone mildly enhances sensitivity to low O2, while co-expression of *UAS-Gyc88E* and *UAS-Gyc89Db* strongly enhances sensitivity to low O2 in A3 v’td1 neurons measured by CaMPARI red/green ratio. Dorsal is top, and anterior is to the left in all images. Scale Bar, 10μm for all images.

**Figure 3 – figure supplement 1.**
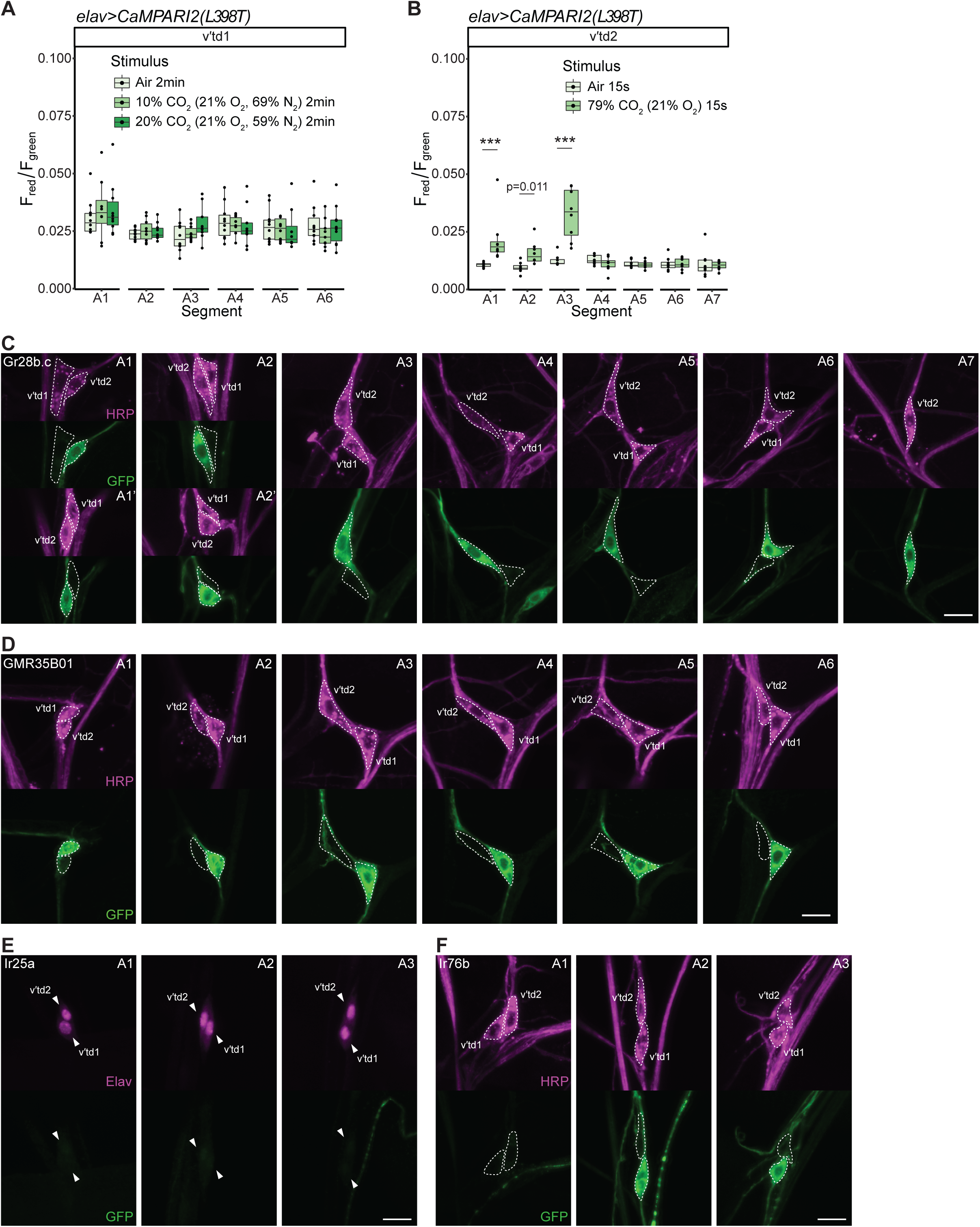
Responses of td neurons to increases in CO2 level and reporter expressions in td neurons. **A.** Quantification of red/green CaMPARI2 fluorescence in v’td1 neurons across segments A1-A6 in response to increased CO2 levels. CaMPARI2 expression was driven by *elav-Gal4*. v’td1 neurons did not show response to increased CO2. n=8-10 animals per condition. **B.** Quantification of red/green CaMPARI2 fluorescence in A1-A7 v’td2 neurons in response to shorter high CO2 exposure. CaMPARI2 expression was driven by *elav-Gal4*. Same three CO2-responding td neurons (A1-A3 v’td2) responded to 15 seconds 79% CO2. n=8 animals per condition. **C.** Images of A1-A7 td neurons in *Gr28b.c-Gal4>UAS-mCD8::GFP* animals. Larval body walls were immunolabeled for horseradish peroxidase (HRP; magenta) and GFP (green). Dorsal is top, and anterior is to the left in all images. *Gr28b.c-Gal4* labels A1-A7 v’td2 neurons and occasionally A1 v’td1 neuron. The position of *Gr28b.c-Gal4*-positive td neurons in A1 segment are occasionally switched along the D-V axis. Same phenomenon has been observed in A2 td neurons. Images of A1 and A2 td neurons show both positions. **D.** Images of A1-A6 td neurons in *GMR35B01-Gal4>UAS-mCD8::GFP* animals. Larval body walls were immunolabeled for horseradish peroxidase (HRP; magenta) and GFP (green). Dorsal is top, and anterior is to the left. *GMR35B01-Gal4* labels A1-A6 v’td1 neurons and occasionally A1 v’td2 neuron. **E.** Images of A1-A3 td neurons in *Ir25a-Gal4> UAS-mCD8::GFP* animals. Larval body walls were immunolabeled for embryonic lethal abnormal vision (Elav; magenta) and GFP (green). Elav labels td nuclei. Dorsal is top, and anterior is to the left. A1-A3 td neurons do not show detectable expression of *Ir25a-Gal4*. **F.** Images of A1-A3 td neurons in *Ir76b-Gal4>UAS-mCD8::GFP* animals. Larval body walls were immunolabeled for horseradish peroxidase (HRP; magenta) and GFP (green). Dorsal is top, and anterior is to the left. A1-A3 v’td2 neurons do not show detectable expression of *Ir76b-Gal4* (Qian et al., 2018).

**Figure 4 – figure supplement 1.**
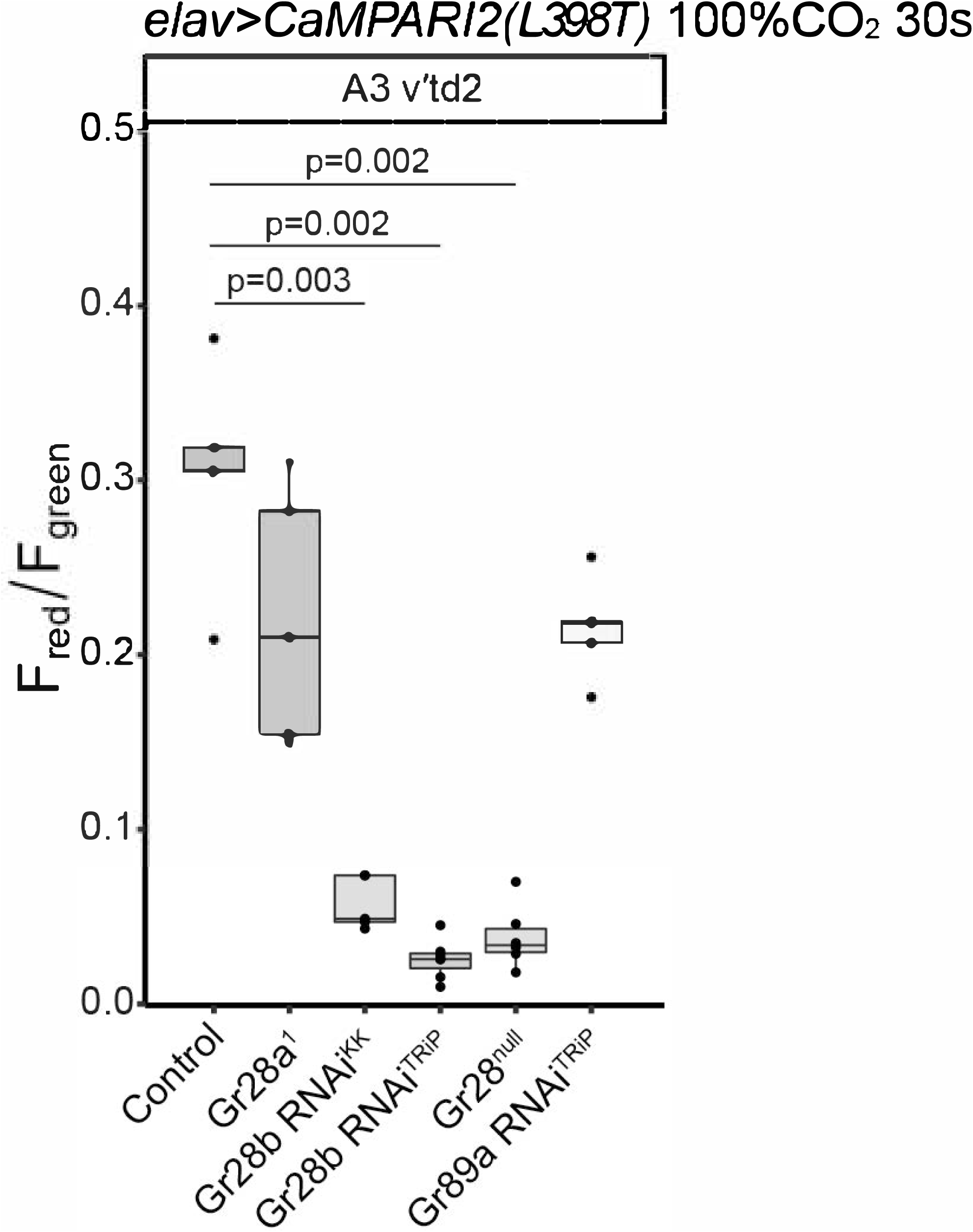
Evidence that Gr28b mediates td neurons responses to CO2. Quantification of red/green CaMPARI2 fluorescence in the CO2-sensitive A3 v’td2 neuron in control and loss-of-function conditions in response to 30 seconds of 100% CO2. RNAi knockdown of Gr28b under the control of *elav-Gal4* and *ΔGr28* significantly decreased the response to high CO2. n=5-7 animals per condition.

**Figure 5 – figure supplement 1.**
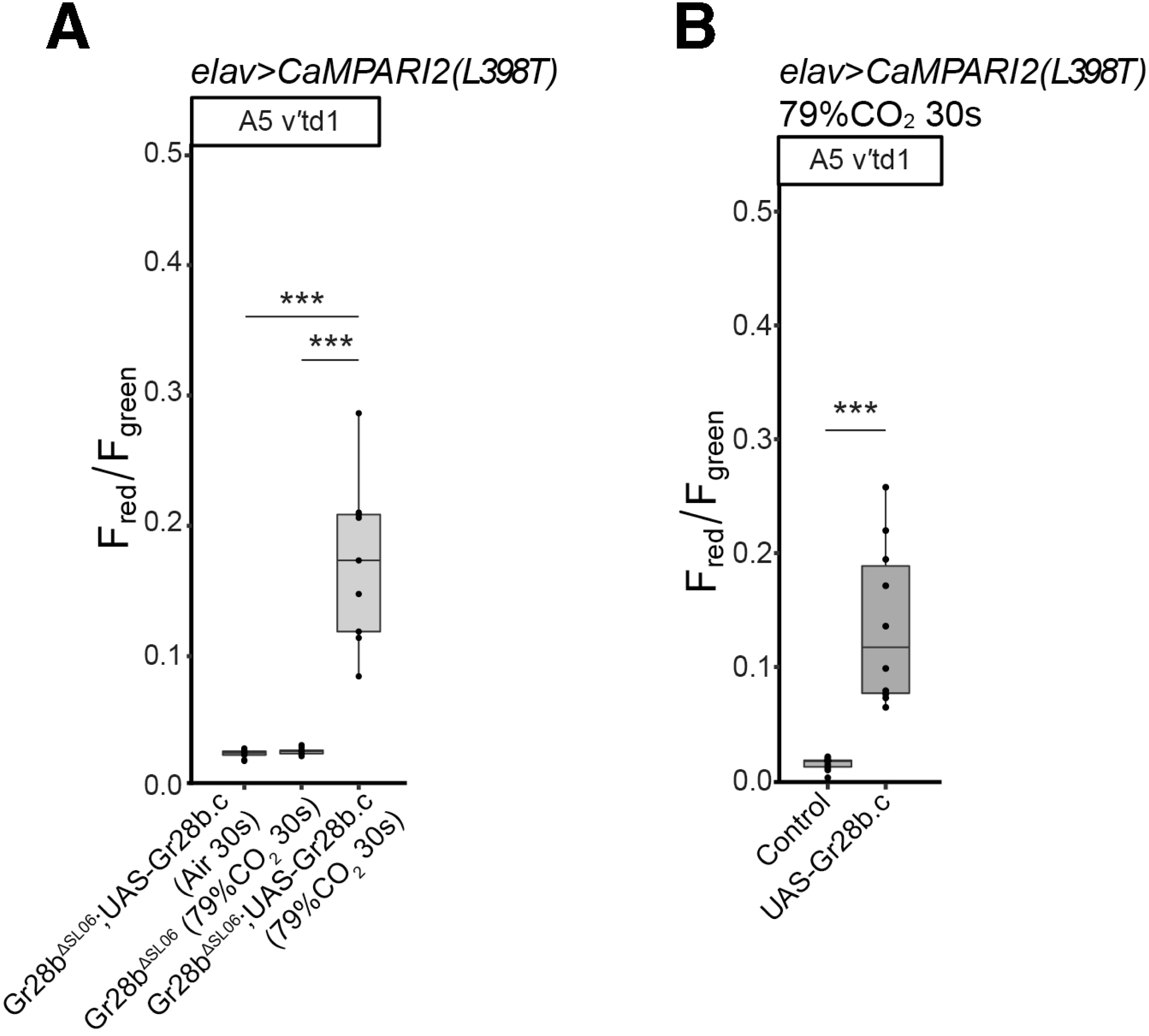
Gr28b.c can induce CO2 sensitivity. **A.** Quantification of red/green CaMPARI2 fluorescence in CO2-insensitive A5 v’td1 neurons in *Gr28b^ΔSL06^*, and *Gr28b^ΔSL06^* with Gr28b.c misexpressed. Larvae mutant for *Gr28b^ΔSL06^*with Gr28b.c misexpressed exposed to 30 seconds air, and *Gr28b^ΔSL06^* larvae exposed to 30 seconds 79% CO2 (same animals as in Figure 5A) showed no td neuron activity. *Gr28b^ΔSL06^* mutants with Gr28b.c misexpressed exposed to 30 seconds 79% CO2 (same animals as in Figure 5A) showed significantly increased activities in v’td1 neurons compared to the other two conditions. n=9 animals per condition. **B.** Quantification of red/green CaMPARI2 fluorescence in CO2-insensitive A5 v’td1 neurons in control and upon ectopic Gr28b.c expression in response to 30 seconds 79% CO2. Ectopic expression of Gr28b.c resulted in significant increases in fluorescence ratio in v’td1 neurons in response to high CO2 level. n=10 animals per condition.

